# A tandem simulation framework for predicting mapping quality

**DOI:** 10.1101/103952

**Authors:** Ben Langmead

## Abstract

Read alignment is the first step in most sequencing data analyses. Because a read’s point of origin can be ambiguous, aligners report a mapping quality: the probability the reported alignment is incorrect. Despite its importance, there is no established and general method for calculating mapping quality. We describe a framework for predicting mapping qualities that works by simulating a set of tandem reads, similar to the input reads in important ways, but for which the true point of origin is known. We implement this in an accurate and low-overhead tool called Qtip, which is compatible with popular aligners.

## Introduction

Read alignment is often the first task when analyzing sequencing data. This is the process of determining each read’s point of origin with respect to a reference genome. Much prior work is concerned with making read aligners computationally efficient [1]. That said, a read’s point of origin can be ambiguous, and the reported alignments can be incorrect [2]. Repetitive genomes, sequencing errors and genetic differences contribute to the problem. In addition to being efficient, aligners must accurately characterize the uncertainty associated with each alignment, as first proposed in the seminal MAQ study [2] which coined the term “mapping quality.” Aligners have methods for predicting mapping quality, which is reported in the MAPQ field of the SAM/BAM format [3]. These methods are generally quite ad hoc, not well described in research literature or software manuals.

We introduce the *tandem simulation* framework for predicting mapping qualities for all the alignments in a dataset in a manner that is agnostic to the aligner and parameters used. We also introduce Qtip, a tool implementing the framework. Qtip operates alongside and in cooperation with an aligner like Bowtie 2 [4]; the term “tandem simulation” refers to this cooperation. After observing the input reads and alignments, Qtip trains an ensemble tree model for predicting mapping qualities. Training uses simulated “tandem” reads, which are randomly drawn from the genome but crafted in a way that mimics statistical properties of the input reads, including their length, quality, gap, and edit distributions. The aligner itself must be modified to report feature data for the model, but alignment algorithms need not be changed. We implemented changes for the Bowtie 2 [4], BWA-MEM [5] and SNAP [6] aligners. Qtip works with any aligner that outputs feature data in a special SAM field; it is not limited to the tools adapted for this study.

We demonstrate Qtip’s predictions are superior to those made by the read aligners themselves, both on average and for most specific MAPQ thresholds tested. We use simulation experiments to show this for various read aligners (Bowtie 2, BWA-MEM, SNAP), alignment settings (read lengths, alignment parameters, species), and accuracy criteria. We also perform a variant-calling experiment to show the improved mapping qualities can benefit downstream analysis. To our knowledge this is the first description of a general technique for characterizing alignment uncertainty, applicable across software tools and alignment settings.

## Background

### Alignment errors

Given a sequencing read and reference genome, a read aligner like Bowtie 2 [4], BWA-MEM [5] or SNAP [6] searches for the read’s highest-scoring alignment to a substring of the reference. Alignment score measures the degree of similarity between the strings, with a higher score indicating fewer mismatches and gaps. If more than one alignment has maximal score, one is chosen arbitrarily. Though many aligners can be configured to report more than one alignment per read, we assume here that just one is reported, as is common. If the reported alignment does not correspond to the read’s true origin, the alignment is *incorrect*, and we call this an *alignment error*. Incorrect alignments lead to interpretation problems later on [7, 8].

Aligners use heuristics — computational shortcuts — to limit effort expended. Heuristics affect which alignments can and cannot be found, shaping what errors the aligner might make. Supplementary Note 1 outlines heuristics used by Bowtie 2.

We can divide alignment errors into three categories, as suggested in the MAQ study [2]:

1. The read is reported to have originated from a locus in the reference genome, but actually originates from a sequence not included in the reference
2. No alignment to the reference is found, but the read actually originates from some locus in the reference
3. An alignment to locus *L*_*r*_ in the reference is reported, but the read actually originates from a different locus in the reference, *L*_*t*_

Category-1 errors might be caused by contaminating DNA, or by an inappropriate or incomplete reference genome sequence. Category-2 errors can occur when the alignment at *L*_*t*_ falls below the minimum similarity threshold (*S*_*min*_), or when the alignment at *L*_*t*_ is missed due to alignment heuristics. Category-3 errors are caused by a combination of repetitive DNA, sequencing errors, genetic differences, and alignment heuristics. Category-3 errors and the related idea of “multi mappers,” reads that align equally well to many loci, are discussed in prior studies [9, 10]. Category-3 errors are also the most numerous, making up 95.8–99.7% of the errors in our simulations (Supplementary Note 2, Supplementary Table 1).

Here we focus on the task of predicting mapping qualities for aligned reads in light of category-3 errors. Category-2 errors are not considered, since no mapping quality prediction is needed in those cases. Although category-1 errors affect mapping quality prediction, we assume they are rare enough to be ignored. In principle, category-1 errors could be included in our model, e.g. by assuming a global prevalence of category-1 error and scaling predictions accordingly, or by including contamination in the simulation.

### Mapping quality

While searching for alignments, aligners uncover information that can be used to predict whether a given alignment is correct. For instance, if the aligner discovers that a read aligns equally well to several copies of a repeat, its confidence that the selected alignment is correct will be low. If the aligner discovers that a read aligns perfectly to one locus and very poorly to a few others, confidence will be higher. Confidence is measured as the probability *p* that the reported alignment is correct. Let the *mapping quality q* = −10 *·* log_10_(1 − *p*). Higher values for *p* (or *q*) indicate higher confidence. The SAM/BAM format [3] requires that *q*, rounded to the nearest integer, be reported in the `MAPQ` field of each alignment. We therefore seek a method that predicts *q* (or equivalently, *p*) accurately across a range of alignment scenarios.

Mapping quality measures something distinct from alignment score. High alignment score indicates high sequence similarity (few mismatches and gaps) between read and reference. It does not imply high mapping quality. For instance, consider a read that aligns with no gaps or mismatches to two distinct loci in the reference. Alignment score is high because there are no gaps or mismatches, but there is only a 50% chance of choosing the correct alignment (*q* ≤ 3). Other measures that do not take genomic repeats into account, such as E-values [11], are also poor proxies for mapping quality.

### Related work

The MAQ study [2] describes sources of alignment error and presents a model for predicting *q* given alignment scores for the best and second-best alignments, and the number of alignments tied for second-best. Successors to MAQ, such as BWA [12], BWA-SW [13] and BWA-MEM [5] use more complex prediction functions. For example, BWA-MEM uses information about whether and how seeds — substrings of the read — match the genome. Qtip uses similar data to train its model. Qtip takes a general approach, learning the prediction model from data, and can adapt to a variety of aligners and alignment settings.

ARDEN [14] uses a mutated “decoy” genome to estimate aggregate prevalence of category-3 errors. However, it is only concerned with aggregate summaries and does not predict *q* for individual alignments. LoQuM [15] uses simulated training alignments and a logistic regression model to predict new *q*s for an already-aligned dataset. Unlike Qtip, LoQuM does not predict *q* from scratch; rather, it “recalibrates” *q* using the aligner-reported mapping quality as an input, along with other inputs derived from the alignment.

The MOSAIK [16] aligner uses a neural network to predict *q*. The user trains the model ahead of time, supplying simulated reads annotated with their true point of origin. Model features include alignment scores of the best and second-best alignments, read sequence entropy, and the number of potential mapping locations. Tandem simulation works similarly to MOSAIK’s approach, building a model from simulated reads, but without requiring the user to collect training data.

Tandem simulation also has similarities to a previous method for allele-specific expression proposed by Hodgkinson et al [17]. In that method, RNA sequencing reads are aligned to a reference genome and allelic ratios are computed at heterozygous sites. The method then simulates a “null” dataset where (a) the genome from which the reads are simulated is customized to include non-reference alleles detected in a separate assay, and (b) when a simulated read overlaps a heterozygous variant, both alleles are sampled with equal frequency. Null reads are aligned to the original reference using the same aligner and parameters as in the initial alignment step, much like the alignment of tandem reads in our framework. Allelic ratios derived from null alignments are used to normalize the original ratios, reducing bias. While our method and Hodgkinson et al’s target different problems, they are alike in their use of a newly simulated dataset to improve results from an initial alignment.

## Results

### Experimental conditions

Simulations were conducted using Mason v0.1.2 [18], or a different simulator where indicated. We ran Qtip v1.6.2 in combination with the Bowtie 2 v2.3.2, BWA-MEM v0.7.15, and SNAP v1.0beta.18. Experiments were performed on nodes of the Maryland Advanced Research Computing Center; each node is an Intel Haswell system with two 12core processors (2.5GHz) and 128GB RAM.

All read aligners were run in their default reporting modes. In other words, all aligners report up to one “best” alignment per read. Reads that fail to align are excluded from the analysis. We used the GRCh38 assembly with some short sequences filtered out (see Supplementary Note 4) as our human reference, except where otherwise noted. Qtip ran on Python v2.7.12 and used scikit-learn v0.18.

### Plots and measures

Let *A* be a vector of *n* alignments *a*_0_*, a*_1_*, …, a*_*n*−1_. Let correct(*a*_*i*_) equal 1 if *a*_*i*_ is correct, 0 otherwise. Let incorrect(*a*_*i*_) equal 1 - correct(*a*_*i*_). An alignment is considered correct if the leftmost base involved in the alignment is within 30 nucleotides of the leftmost base in the simulated substring, with appropriate adjustments for soft clipping. Let *Q* = *q*_0_*, q*_1_*, …, q*_*n*−1_ be mapping qualities corresponding to *a*_0_*, a*_1_*, …, a*_*n*−1_, as predicted by the read aligner, and let *P* = *p*_0_*, p*_1_*, …, p*_*n*−1_ be the corresponding correctness probabilities, using the relationship that *q* = −10 *·* log_10_(1 − *p*). Let *Q′* and *P ′* be defined similarly, but for the mapping qualities predicted by Qtip.

We define plots (CID, CSED) and measures (RCA, RCE) that characterize how Qtip’s predictions (*Q′*) compare to the aligner’s (*Q*). CID and RCA capture how well *Q′ ranks* alignments from most to least likely to be correct relative to *Q*. CID and RCA are invariant under monotonic transformations of *P* and *P ′*; they are concerned only with how well alignments are ranked, not with probabilities per se. CSED and RCE capture how closely *P′* matches the the true correctness relative to *P* ; i.e. CSED and RCE are concerned with how well *P ′* and *P* fit their probabilistic interpretation.

### Cumulative Incorrect Difference (CID)

Let *Â*be *A* sorted in descending order by *Q*, and likewise for *Â′* and *Q′*. The *cumulative incorrect vector C* is the vector *c*_0_*, c*_1_*, …, c*_*n*−1_ such that 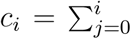 incorrect 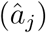. *C′* is defined similarly for *Â′*. Let *D* be the element-wise difference *C′ -C*. When *d*_*i*_ *<* 0, Qtip’s mapping qualities yield a better segregation of correct from incorrect alignments about the “pivot” *i*. When *d*_*i*_ > 0, the aligner’s mapping qualities give the better segregation. A “CID plot” draws a line representing the *d*_*i*_’s (vertical axis) for *i* = 0 to *n* − 1 (horizontal axis), and we judge Qtip’s efficacy according to the line’s tendency to stay below *y* = 0.

### Cumulative Squared Error Difference (CSED)

Let *Â*and 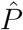 be *A* and *P* sorted in descending order by *P*, and likewise for *Â ′* and 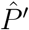. The *cumulative squared error vector E* is the vector *e*_0_*, e*_1_*, …, e*_*n*−1_ such that 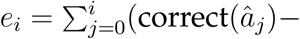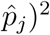, with *E′* defined similarly for *Â ′* and *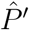*^2^*. Let S* be the element-wise difference *E′ -E*. When *s*_*i*_ *<* 0, Qtip’s mapping qualities yield lower squared error up to the *i*th alignment. The “CSED plot” draws a line representing the *s*_*i*_’s (vertical axis) for *i* = 0 to *n* −1 (horizontal axis). Like for the CID plot, we judge Qtip’s efficacy according to the line’s tendency to stay below *y* = 0.

### Relative change in area under CID (RCA)

is defined as 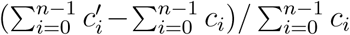. Negative values indicate a better overall ranking is achieved using Qtip’s predictions.

### Relative change in sum of squared errors (RCE)

is defined as (SSE(*P ′*) - SSE(*P*))/SSE(*P*), where 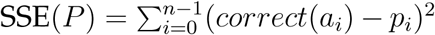. Negative values indicate that Qtip’s predictions yield lower total squared error.

The distinction between the rank-based (CID, RCA) and probabilistic (CSED, RCE) metrics relates to how downstream tools, e.g. variant callers, use mapping qualities. Freebayes [19] and the Genome Analysis Toolkit (GATK) [20] ignore an alignment if its mapping quality is below a threshold. In this case, CID and RCA are relevant since as directly evaluate how well various thresholds separate correct from incorrect alignments. Other methods, such as the consensus genotype calling method described in the MAQ study [2], interpret a mapping quality as a probability. Alignments are weighted according to their probability, with no alignments excluded. Here, CSED and RCE are relevant since they directly evaluate how well the probabilities match actual correctness status.

We note that the problem of evaluating and plotting relative quality of two sets of mapping-quality predictions is not specifically addressed in past studies. ROC-like plots are used for the related task of comparing aligners [4, 5], where the axes represent false and true positives and a line follows points corresponding to increasingly permissive mapping-quality thresholds. But the two-dimensionality of these plots makes it hard to find comparable points, points on two curves where the threshold allows same number of alignments. A similar problem exists for comparisons examining particular thresholds (*≥* 10, ≥ 20, etc); for two sets of predictions the thresholds might allow very different numbers of alignments, impeding interpretation. CID and CSED plots are inspired by Accuracy versus Drop Rate (ADR) plots [21] and are related to ROC-like plots, except: (a) two lines are represented more concisely as a single line giving the difference, and (b) at a given horizontal point, we are comparing thresholds that allow the same number of alignments (the same “drop rate”).

## Simulation experiments

We conducted simulation experiments to show how Qtip’s mapping quality predictions compare to those made by read aligners themselves. We vary several experimental conditions, including (a) read length, (b) aligner parameterization, (c) reference genome, (d) read alignment tool, and (e) read simulator. The simulator encodes the read’s true point of origin in the read name, allowing Qtip to check later whether an alignment is correct.

### Simulated samples

We used Mason to simulate five Illumina-like samples with unpaired reads of length 50, 100, 150, 250 and 500 respectively. We simulated five paired-end samples with the same lengths, with most fragment lengths between 2*L* − 4*L* nt where *L* is the read length. We simulated 4 million reads/pairs for each sample. Aligners were configured to consider fragment lengths in the 2*L* − 4*L* range as concordant. Thus, most simulated pairs aligned concordantly (consistent with paired-end constraints) whereas some aligned discordantly. Simulator commands, and implications for fragment lengths, are discussed in Supplementary Note 4. Alignment commands are in Supplementary Note 5.

### Varying read length

We used Qtip together with Bowtie 2 to align and predict mapping qualities for each Mason-simulated sample. We rounded Qtip’s predictions to the nearest integer per the SAM/BAM format. For each alignment, we parsed aligner-predicted and Qtip-predicted mapping qualities, as well as the read’s true point of origin as provided by Mason. We calculated RCA and RCE (Table 2) and plotted CSED (Figure 1). CSED *y* values were scaled with *y*_*plot*_ = sign(*y*_*orig*_) *·* log_10_(*|y*_*orig*_*|* + 1).

**Figure 1:**
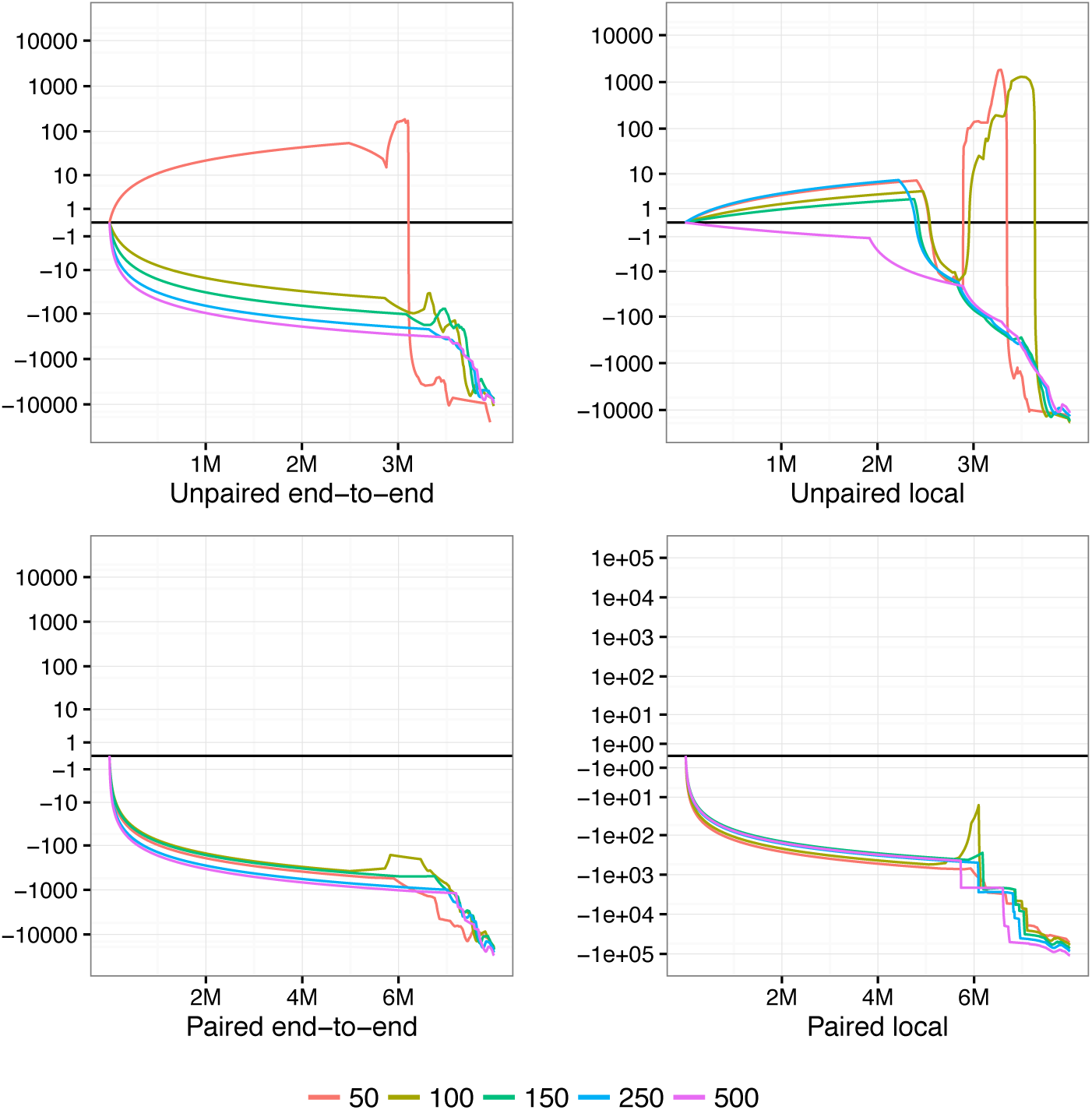
CSED for various lengths. Cumulative squared-error difference plot from running Qtip and Bowtie 2 on Mason-simulated Illumina-like samples of various lengths. Each sample consists of 4 million reads or pairs. The horizontal axis indicates cumulative number of reads/ends passing the threshold, with the left-hand extreme corresponding to a high mapping-quality threshold and right-hand extreme corresponding to a low threshold. Results for unpaired samples are on top, paired on bottom. Bowtie 2 is run in its (default) end-to-end alignment mode in the case of the left-hand plots, and in local alignment mode in the case of the right-hand plots.

To measure variability of Qtip’s predictions, we repeated each experiment 10 times starting from step 2 onward, seeding the pseudo-random number generator differently in each trial. RCA/RCE tables describe all 10 trials whereas, for clarity, the CSED plot describes only the first trial.

Qtip’s mapping qualities are overall superior to those predicted by Bowtie 2, as indicated by the negative RCAs and RCEs (Table 2). This is true across all samples tested, and in both end-to-end and local alignment mode. The improvement is larger for samples with longer reads and for paired-end samples. Variability is modest overall but somewhat higher for longer reads. See Discussion for further comments on variability.

There are portions of the CSED plots (Figure 1) where the plot rises above *y* = 0, indicating the aligner-reported mapping qualities exhibit lower cumulative squared error at those thresholds. This is most prominent in the unpaired experiments, particularly for 50 nt reads. However, Qtip’s superior predictions at other *q* thresholds – especially low ones – help bring overall RCE below zero in all cases. For paired-end samples, CSEDs show Qtip’s predictions are superior at nearly all *q* thresholds.

### Varying reference genome and alignment tool

To study how genomes of varying length and repetitiveness influence Qtip’s performance, we experimented with four reference genome assemblies spanning three species: Human GRCh37, Human GRCh38, Mouse GRCm38, and *Zea mays* AGPv4. The human GRCh38 primary assembly is 3.10 Gbp long (2.95 Gbp excluding Ns) with 50% of the genome annotated as repetitive according to RepeatMasker [22]. GRCh37 is 3.10 Gbp long (2.86 Gbp excluding Ns) with 47% of the genome annotated as repetitive. GRCm38 is 2.73 Gbp long (2.65 Gbp excluding Ns), with 44% of the genome annotated as repetitive. AGPv4 is 2.13 Gbps long (2.10 Gbp excluding Ns). Though no official RepeatMasker annotation is available, past studies report that 85% of the genome consists of transposable element sequences [23], making it the most repetitive of the genomes tested. We used the Mason-simulated 100 nt and 250 nt samples, both unpaired and paired-end. We tested three aligners – Bowtie 2, BWA-MEM and SNAP– with each genome. The changes made to each aligner in order to work with Qtip are detailed in Supplementary Note 6. We calculated RCA and RCE for the 10 trials and plotted CSED for only the first trial.

**Table 1:** Relative change in area under CID (RCA) and relative change in sum of squared error (RCE) when running Qtip and Bowtie 2 on Mason-simulated Illumina-like samples of various lengths. Relative change is expressed as a percent. Each sample consists of 4 million reads/pairs. Samples are either unpaired and paired-end, and Bowtie 2 is run in either end-to-end or local alignment mode as indicated. Results are means and standard deviations over 10 random trials, repeated starting from the input modeling step.

|  |  | End-to-end |  |  |  | Local |  |  |  |
| --- | --- | --- | --- | --- | --- | --- | --- | --- | --- |
|  |  | RCA |  | RCE |  | RCA |  | RCE |  |
|  | Read length | mean | sd | mean | sd | mean | sd | mean | sd |
| Unpaired | 50 | -9.03 | 0.26 | -24.61 | 0.19 | -2.49 | 0.47 | -15.30 | 0.57 |
|  | 100 | -7.43 | 1.82 | -18.53 | 2.61 | -10.26 | 1.52 | -27.87 | 1.59 |
|  | 150 | -9.11 | 1.15 | -16.22 | 0.62 | -14.77 | 2.15 | -29.27 | 1.03 |
|  | 250 | -16.82 | 2.01 | -19.97 | 0.44 | -22.51 | 1.77 | -28.85 | 0.62 |
|  | 500 | -33.77 | 0.60 | -27.26 | 0.37 | -37.75 | 1.32 | -31.65 | 1.01 |
| Paired | 50 | -13.11 | 0.44 | -19.60 | 0.44 | -18.60 | 0.28 | -33.63 | 0.24 |
|  | 100 | -15.80 | 1.29 | -21.79 | 0.33 | -36.84 | 0.69 | -45.42 | 0.53 |
|  | 150 | -22.39 | 1.74 | -25.94 | 0.28 | -46.87 | 0.54 | -52.76 | 0.56 |
|  | 250 | -37.65 | 0.97 | -33.48 | 0.22 | -58.39 | 0.91 | -58.75 | 1.38 |
|  | 500 | -54.59 | 0.58 | -44.91 | 0.34 | -68.54 | 0.95 | -70.37 | 3.45 |

Qtip-predicted mapping qualities are superior in nearly all scenarios, as indicated by negative RCAs and RCEs (Table 2). The exceptions are three of the human paired-end SNAP experiments (GRCh37 100 nt, GRCh37 250 nt and GRCh38 250 nt), which have negative RCA but positive RCE. Variability of RCAs and RCEs across trials is generally modest, but tool dependent, with SNAP exhibiting the highest variabilities. BWA-MEM’s standard deviations are small, all below 0.6. Bowtie 2’s range up to 2.61 and SNAP’s up to 4.44. See Discussion for further comments on variability.

CSED curves (Figure 2) again show that for some thresholds, aligner-reported mapping qualities are superior in terms of minimizing cumulative squared error, i.e. where the CSED rises above *y* = 0. Qtip’s mapping qualities seem to perform worse for many thresholds in the BWA-MEM unpaired experiments, especially for *Zea mays*. But Qtip’s qualities consistently perform better at very low thresholds. Qtip’s mapping qualities perform particularly well for the Bowtie 2 *Zea mays* experiments, and for all the paired-end experiments.

**Figure 2:**
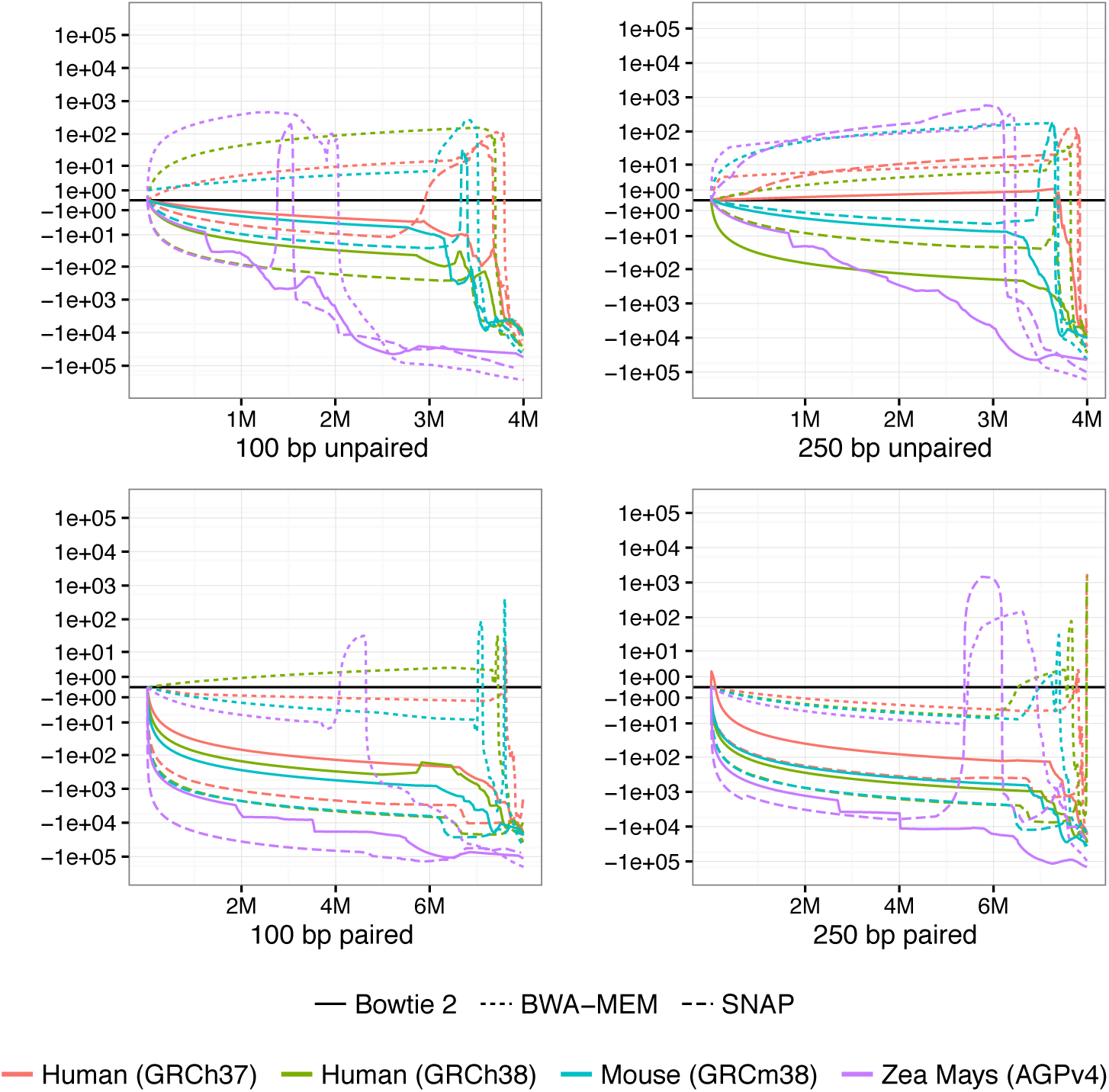
CSED for various aligners and references. Cumulative squared-error difference plot from running Qtip with various reference genomes and read aligners. The input reads are Mason-simulated Illumina-like 100 nt (left) and 250 nt (right) samples, each consisting of 4 million reads/pairs. The horizontal axis indicates cumulative number of reads/ends passing the threshold, with the left-hand extreme corresponding to a high mapping-quality threshold and right-hand extreme corresponding to a low threshold. Results for unpaired samples are shown on top, and paired on bottom.

**Table 2:** Relative change in area under CID (RCA) and relative change in sum of squared error (RCE) for various aligners and reference genomes, expressed as percent change. The experiments used 100 nt or 250 nt reads, and unpaired or paired-end reads, as indicated. Results are means and standard deviations over 10 random trials, repeated starting from the input modeling step.

|  |  |  | 100 nt |  |  |  | 250 nt |  |  |  |
| --- | --- | --- | --- | --- | --- | --- | --- | --- | --- | --- |
|  |  |  | RCA |  | RCE |  | RCA |  | RCE |  |
|  |  |  | mean | sd | mean | sd | mean | sd | mean | sd |
| Unpaired | GRCh37 | Bowtie 2 | -11.22 | 1.08 | -24.43 | 0.66 | -15.02 | 0.39 | -28.37 | 0.38 |
|  |  | BWA-MEM | -14.49 | 2.29 | -49.54 | 0.43 | -9.14 | 2.09 | -52.31 | 0.38 |
|  |  | SNAP | -15.94 | 0.32 | -36.86 | 0.23 | -9.53 | 3.88 | -28.57 | 0.47 |
|  | GRCh38 | Bowtie 2 | -7.43 | 1.82 | -18.53 | 2.61 | -16.82 | 2.01 | -19.97 | 0.44 |
|  |  | BWA-MEM | -15.49 | 0.58 | -47.42 | 0.37 | -15.78 | 0.57 | -51.14 | 0.31 |
|  |  | SNAP | -19.58 | 0.18 | -36.58 | 0.47 | -14.74 | 0.27 | -25.47 | 0.32 |
|  | Mouse | Bowtie 2 | -5.60 | 0.24 | -17.19 | 0.45 | -7.05 | 0.33 | -17.73 | 0.37 |
|  |  | BWA-MEM | -13.50 | 0.15 | -46.25 | 0.27 | -16.39 | 0.38 | -51.12 | 0.30 |
|  |  | SNAP | -9.02 | 0.17 | -31.07 | 0.33 | -10.61 | 0.20 | -31.78 | 0.43 |
|  | Zea mays | Bowtie 2 | -6.63 | 0.32 | -19.56 | 0.25 | -17.09 | 0.38 | -25.89 | 0.44 |
|  |  | BWA-MEM | -19.26 | 0.11 | -58.32 | 0.26 | -25.14 | 0.19 | -66.76 | 0.23 |
|  |  | SNAP | -13.02 | 0.24 | -38.48 | 0.53 | -24.01 | 0.43 | -53.80 | 0.42 |
| Paired | GRCh37 | Bowtie 2 | -25.86 | 0.33 | -30.26 | 0.50 | -36.31 | 2.16 | -38.29 | 0.60 |
|  |  | BWA-MEM | -13.33 | 0.23 | -45.70 | 0.27 | -10.08 | 0.75 | -47.58 | 0.31 |
|  |  | SNAP | -56.53 | 1.99 | 1.39 | 2.34 | -42.89 | 7.63 | 13.17 | 3.95 |
|  | GRCh38 | Bowtie 2 | -15.80 | 1.29 | -21.79 | 0.33 | -38.22 | 0.31 | -33.63 | 0.30 |
|  |  | BWA-MEM | -14.19 | 0.16 | -41.35 | 0.19 | -12.36 | 0.46 | -42.78 | 0.29 |
|  |  | SNAP | -51.36 | 0.98 | -11.16 | 1.12 | -51.32 | 1.45 | 4.34 | 2.29 |
|  | Mouse | Bowtie 2 | -10.10 | 0.26 | -18.93 | 0.31 | -19.03 | 0.21 | -29.07 | 0.32 |
|  |  | BWA-MEM | -11.86 | 0.12 | -36.18 | 0.37 | -13.30 | 0.19 | -39.91 | 0.21 |
|  |  | SNAP | -29.90 | 0.67 | -17.04 | 0.76 | -30.16 | 0.30 | -15.79 | 0.47 |
|  | Zea mays | Bowtie 2 | -17.92 | 0.21 | -26.95 | 0.27 | -43.19 | 0.18 | -51.69 | 0.38 |
|  |  | BWA-MEM | -17.04 | 0.15 | -47.48 | 0.29 | -21.45 | 0.20 | -56.58 | 0.08 |
|  |  | SNAP | -36.28 | 0.55 | -17.08 | 0.79 | -26.45 | 4.44 | -20.05 | 0.52 |

To assess how greater incidence of Category-1 errors affects results, we repeated the human experiments, expanding the simulation to include reads both from the reference genome and from sequences in the CHM1 hydatidiform mole assembly not present in the reference. We used Assemblytics [24] to obtain CHM1-specific sequences as detailed in Supplementary Note 3. The results show this has little effect on the accuracy of Qtip’s predictions (Supplementary Table 2).

### Other simulation experiments

We also conducted simulation experiments varying the sensitivity level of the aligner, described in Supplementary Note 7, Supplementary Table 3 and Supplementary Figure 1, and varying the software tool used to generate the simulated reads, described in Supplementary Note 8, Supplementary Table 4 and Supplementary Figure 2.

### Variant calling

To demonstrate Qtip’s effect on downstream results, we evaluated variant calling accuracy with and without Qtip’s predictions. We used paired-end human 100 x 100 Illumina HiSeq reads from the Platinum Genomes project [25] (ERR194147) and gold-standard Platinum variants [25] for the sequenced individual (NA12878). The Platinum variants are high-confidence pedigree-validated calls supported by multiple bioinformatics pipelines and sequencing technologies. The analysis is limited to areas of the genome called with high confidence by Platinum Genomes.

We used Freebayes v1.1.0 [19] to call single-nucleotide variants (SNVs) once for the alignments with the original mapping qualities and again for the same alignments but with Qtip-predicted mapping qualities. Following past studies [26], we filtered out variant calls with read depth greater than 4 poisson standard deviations above the mean. We defined a true positive as an SNV call made from ERR194147 data that matched a platinum call, a false positive as a call made from ERR194147 that did not match any platinum call, and a false negative as a platinum call that did not match any ERR194147 call. We calculated *F*_*β*_ for various *β*s. *F*_1_ (*β* = 1) is the typical F1-score, related to the harmonic mean of precision and recall. Setting *β >* 1 gives recall more weight than precision and setting *β* < 1 gives precision more weight than recall. We tried values of *β* ranging from 0.25 to 4 to cover a range of precision-recall tradeoffs. Further details on alignment and variant calling are given in Supplementary Note 9.

Like other variant callers and downstream tools, Freebayes thresholds on mapping quality (*Q*) to eliminate some alignments prior to variant calling, eliminating alignments with *Q* < 1 by default. Since we are concerned with overall accuracy of mapping qualities and not with any particular threshold, we re-ran Freebayes with various integer *Q* thresholds: 0–12, 15, 20 and 30. Freebayes also associates a genotype quality value with each called variant, given in the VCF file’s `QUAL` field. We used the `vcfroc` tool from `vcflib` (https://github.com/vcflib/vcflib) to evaluate all possible `QUAL` thresholds for all possible *Q* thresholds, ultimately selecting *Q* and `QUAL` thresholds maximizing *F*_*β*_.

**Table 3:** Single-nucleotide variant (SNV) *F*_*β*_ scores for various *β*s with original mapping qualities and with Qtip-generated qualities. Paired-end reads from ERR194147, a female, were aligned with Bowtie 2 together with Qtip. SNV variants were called with Freebayes for chromosomes 1–22 and X. Variant-quality (QUAL) and mapping-quality (Q) thresholds yielding greatest *F*_*β*_ score are reported. Platinum variants were used as the true callset. Before calculating *F*_*β*_, calls outside Platinum Genomes high-confidence regions were excluded. Rightmost columns show differences in *F*_*β*_, number of true positive SNVs, and number of false positive SNVs.

| $\beta$ | Original | | | Qtip | | | $\Delta$ (Qtip – Orig) | | |
| --- | --- | --- | --- | --- | --- | --- | --- | --- | --- |
| | $F_\beta$ | QUAL | Q | $F_\beta$ | QUAL | Q | $F_\beta$ | TP | FP |
| 0.250 | 0.9925 | 194 | 2 | 0.9924 | 213 | 2 | -2.2e-05 | +20,189 | +1,505 |
| 0.333 | 0.9906 | 150 | 3 | 0.9908 | 178 | 2 | +2.6e-04 | +16,001 | +911 |
| 0.500 | 0.9872 | 87 | 3 | 0.9881 | 125 | 3 | +8.3e-04 | +15,143 | +311 |
| 0.750 | 0.9843 | 10.6 | 3 | 0.9854 | 70.1 | 4 | +1.1e-03 | -413 | -6,537 |
| 1.000 | 0.9835 | 0.0158 | 3 | 0.9845 | 13.6 | 4 | +9.4e-04 | +3,999 | -2,745 |
| 1.500 | 0.9846 | 1.79e-06 | 3 | 0.9856 | 0.000675 | 5 | +1.0e-03 | +4,392 | -1,832 |
| 2.000 | 0.9860 | 5.06e-08 | 3 | 0.9870 | 1.38e-05 | 5 | +1.0e-03 | +3,110 | -5,692 |
| 3.000 | 0.9880 | 1.16e-09 | 3 | 0.9889 | 8.64e-08 | 4 | +8.6e-04 | +2,583 | -7,937 |
| 4.000 | 0.9892 | 1.06e-10 | 3 | 0.9899 | 6.58e-09 | 4 | +7.1e-04 | +1,891 | -13,600 |

Results are presented in Table 2. For all *β*s examined except the lowest (*β*=0.25), Qtip-predicted mapping qualities yielded superior *F*_*β*_. For *β* ≥ 1, Qtip’s predictions yielded more true positives and fewer false positives than the original predictions. For 0.25 ≤ *β ≤* 0.5 Qtip’s predictions yielded around 15,000–20,000 more true positives at the cost of around 300–1,500 more false positives. Notably, these improvements were achieved simply by changing the mapping qualities; the alignments are the same and the variant caller has not been modified or tuned in any way. We also note that Qtip’s improved performance is obtained using a smaller range of mapping quality values. Qtip-predicted mapping qualities in this experiment ranged from 0 to 36, whereas Bowtie 2 mapping qualities ranged from 0 to 42.

### Efficiency and overhead

The tandem simulation framework adds overhead to the alignment process. We measured Qtip’s overhead when analyzing public datasets ERR050082 & ERR050083. Specifically, we measured how running time and peak memory footprint grew when Qtip ran alongside the aligner, versus when the aligner ran by itself. Running-time overhead is modest for Bowtie 2 and BWA-MEM, ranging from 5% to 10% (Table 4). For SNAP, running-time overhead is larger, 12% to 14% for unpaired and 23% to 28% for paired-end alignment. Peak memory footprint added by Qtip was 200-400 megabytes in all cases, substantially smaller than the footprint of the aligners themselves which must keep a copy of the reference genome index in memory. In the case of SNAP, peak memory footprint increased by less than 1.15%. For BWA-MEM the increase was always less than 5% and for Bowtie 2 less than 13%.

## Methods

### Tandem simulation

The user specifies a collection of input reads: *R* = *r*_0_*, r*_1_*, …, r*_*n*−1_, a read aligner, alignment parameters, a reference genome in FASTA format, and any other files required, such as a genome index. The tandem simulation framework (Figure 3) aligns the input reads to the reference genome and predicts a mapping quality *q*_*i*_ for each aligned read. In step 1, input reads are aligned to the reference genome using the specified aligner and parameters. In step 2, the SAM-formatted [3] alignments are parsed and an *input model*, capturing information about the input reads and their alignments, is built. In step 3, the input model and reference genome are used to simulate a new set of reads, called *tandem* reads since they originate from tandem simulation. Each tandem read is from a random location in the genome and is labeled with its true point of origin. In step 4, tandem reads are aligned to the reference genome using the same aligner and parameters as in step 1. In step 5, the alignments produced in step 4 are parsed and converted to *training records*. Because the true point of origin is known, each training record can be labeled as correct or incorrect. In step 6, a model is trained on the records from step 5. In step 7, SAM alignments from step 1 are parsed. For each aligned read, a *test record*, like the training record from step 5, is constructed. Based on the test record, the trained model is applied to predict *q*_*i*_. The alignment’s SAM record is then re-written substituting *q*_*i*_ in the MAPQ field. New predictions for all input alignments are written in this way.

**Figure 3:**
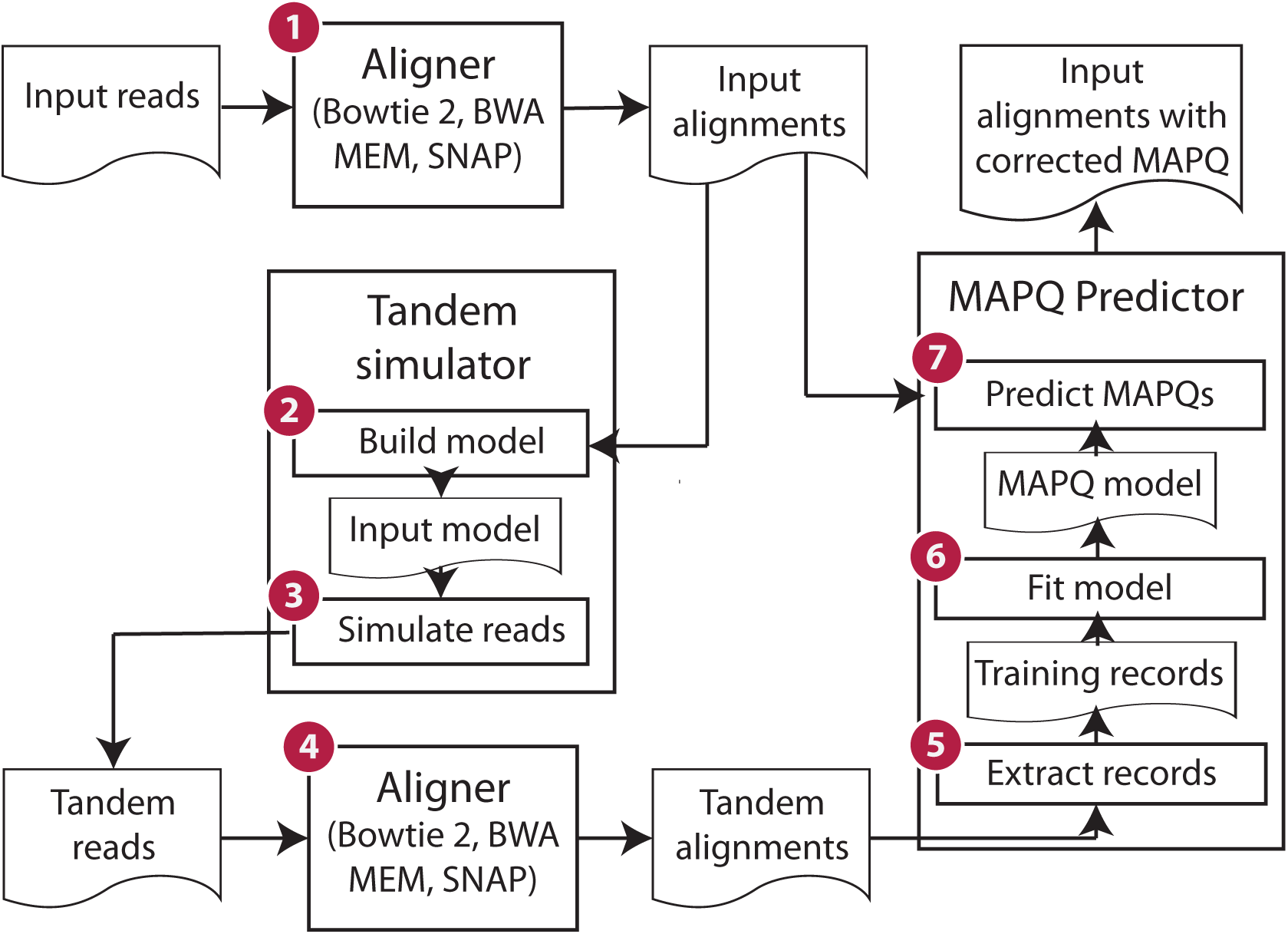
Stages of Qtip pipeline Computational steps and intermediate results in Qtip. Numbers denote ordering of steps and arrows denote flow of data. The input (upper left) is a collection of sequencing reads and the ultimate output (upper right) is SAM file containing alignments, where each aligned read’s MAPQ field is set according to Qtip’s prediction.

**Table 4:** Overhead of the Qtip tool. Measured as the increase in running time (left) and peak memory footprint (right) from when the aligner runs by itself (“Time”) to when the aligner runs in combination with Qtip (“+Qtip”). “% inc” columns give the percent increase. Times are in minutes and memory footprints are in gigabytes.

|  |  |  | Time (minutes) |  |  | Peak memory (gigabytes) |  |  |
| --- | --- | --- | --- | --- | --- | --- | --- | --- |
|  |  |  | Time | +Qtip | % inc | Memory | +Qtip | % inc |
| ERR050082 | Unpaired | Bowtie 2 | 23.58 | 25.20 | 6.89 | 3.26 | 3.52 | 8.25 |
|  |  | BWA-MEM | 22.18 | 23.75 | 7.08 | 7.53 | 7.79 | 3.40 |
|  |  | SNAP | 12.13 | 13.75 | 13.32 | 29.26 | 29.51 | 0.85 |
|  | Paired | Bowtie 2 | 57.52 | 61.35 | 6.67 | 3.27 | 3.66 | 12.02 |
|  |  | BWA-MEM | 57.93 | 63.38 | 9.41 | 7.87 | 8.26 | 4.87 |
|  |  | SNAP | 11.28 | 14.42 | 27.69 | 30.21 | 30.55 | 1.14 |
| ERR050083 | Unpaired | Bowtie 2 | 23.02 | 24.75 | 7.55 | 3.26 | 3.53 | 8.28 |
|  |  | BWA-MEM | 24.60 | 26.08 | 6.03 | 7.75 | 8.01 | 3.33 |
|  |  | SNAP | 12.23 | 13.73 | 12.31 | 29.26 | 29.51 | 0.85 |
|  | Paired | Bowtie 2 | 63.60 | 67.40 | 6.00 | 3.27 | 3.66 | 11.96 |
|  |  | BWA-MEM | 61.58 | 67.52 | 9.62 | 7.92 | 8.31 | 4.83 |
|  |  | SNAP | 11.95 | 14.68 | 22.86 | 30.21 | 30.55 | 1.14 |

Importantly, the mapping quality model trained in step 6 is tailored to the alignment scenario at hand. The aligner and parameters from step 1 are reused in step 4, and tandem reads generated in step 3 mimic the input reads.

To work with the tandem simulation framework, the aligner must report feature data — how well the read aligned, what other alignments were found, what heuristics were used, etc — used to train and apply the model. This requires modifications to the alignment software. The modifications are not complex and do not affect efficiency or accuracy of the aligner. However, making appropriate modifications requires knowledge of how the aligner works and of which intermediate alignment results constitute informative features. For this study, we adapted three tools: Bowtie 2 v2.3.2, BWA-MEM v0.7.15 and SNAP v1.0beta.18. Supplementary Note 6 provides links to our modifications and details about the modifications made and how features were chosen.

We chose these three aligners both because of their popularity and because they together support a breadth of alignment scenarios. For example, Bowtie 2 and BWA-MEM support local alignment, Bowtie 2 and SNAP support end-to-end alignment, and all three tools support both unpaired and paired-end alignment. Also, all three tools produce their own mapping quality predictions. We will evaluate Qtip’s mapping quality predictions in all of these modes, comparing Qtip’s predictions to those made by the aligners.

### Read and alignment categories

When predicting mapping quality, Qtip uses a different model depending on whether the alignment is unpaired (*unp*), paired-end and concordantly aligned (*conc*), paired-end and discordantly aligned (*disc*), or paired-end with the opposite end having failed to align (*bad-end*). Qtip trains each model with alignments of the same category. Qtip parameters control the minimum number of tandem reads or pairs of each category to generate. The default number for each category is *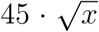* where *x* is the number of input alignments of that category. Both the scaling factor and the function are configurable via Qtip’s `--sim-function` and `--sim-factor` parameters. Qtip enforces a minimum of 30,000 tandem reads in the case of the *conc* and *unp* categories and 10,000 in the case of the *disc* and *bad-end* categories. The formula for number of training records is discussed further in Supplementary Note 10, with alternatives explored in Supplementary Figure 3.

### Input model and simulation of tandem reads

The input model built in step 2 of Qtip (Figure 3) captures information about the input reads and alignments. Qtip uses this to simulate new tandem reads that are from random genomic locations but are similar to the input reads in key ways, mimicking their read length distribution, quality strings, and patterns of gaps and mismatches. Tandem paired-end reads additionally mimic the input’s fragment length distribution and relative orientation of the two ends.

To accomplish this, Qtip takes the following approach. For each aligned unpaired read, a *template* data record is created. The template consists of the strand aligned to, the read’s quality string, and the pattern of mismatches and gaps in the alignment as defined by the `CIGAR` and `MD:Z` SAM fields. For each aligned pair, the template additionally stores the pair’s inferred fragment length and a flag indicating which end aligned upstream with respect to the genome. Since templates for large datasets can quickly exhaust memory, Qtip uses reservoir sampling keep a configurable-sized subsample of the templates. The default sample size is 10,000.

In step 3, Qtip uses the input model to simulate tandem reads. To simulate an unpaired tandem read, Qtip randomly draws an unpaired template, with replacement and uniform probability, from those collected in step 2. A new read is constructed from the template by (a) drawing an appropriate-length substring from the reference genome uniformly at random, (b) possibly reverse-complementing it, according to the template strand, (c) mutating the extracted sequence according to the template pattern of mismatches and gaps, and (d) setting the new read’s quality string equal to the template’s. The simulated read’s point of origin is encoded in the read name, allowing later steps to check whether an alignment is correct. The process for simulating a paired tandem read is similar, with fragment length determined by the template. More details are given in Supplementary Note 11.

Importantly, some aspects of the input data are hard to mimic. For example, errors made by 454 and Ion Torrent sequencing technologies can manifest as spurious extensions or retractions of homopolymers. Since genome substrings are matched with templates randomly, homopolymer errors in the template will often fail to line up with homopolymers in the substring. Other aspects of the input data are not as difficult to mimic, but happen not to be captured in Qtip’s simulation. For example, if a dataset is enriched or depleted for reads drawn from a particular genomic feature (e.g., coding regions), Qtip’s simulation, which draws reads uniformly at random from across the genome, will not exhibit that pattern. While we demonstrate Qtip performs well despite these deficiencies, they nonetheless illustrate that it is difficult to construct tandem reads that truly mimic input reads in all ways. We return to this in the Discussion section.

### Mapping quality model

Given training records derived from tandem reads aligned in step 4, we train a model in steps 5 and 6 that is later used to predict *q*s for the input alignments. Qtip trains separate models for each alignment category: *unp*, *conc*, *disc*, and *bad-end*. The particular features used to train a model vary depending on the alignment category and read aligner. We briefly summarize them here, but more details are provided in Supplementary Note 6.

These features are included regardless of aligner or alignment category: (a) the alignment score of the best alignment, (b) the difference between the alignment score of the best alignment and that of the second-best alignment if one was found, (c) the length of the aligned read, (d) the sum of the base qualities of the aligned bases, and (e) the sum of the base qualities of the soft-clipped bases. In the case of a concordantly-aligned paired, the inferred fragment length (from the SAM TLEN field) is also included as a feature.

By default Qtip uses an implementation of random forests [27] from the scikit-learn [28] library to model and predict mapping qualities. The random forest consists of many decision trees, each trained on a bootstrap sample of the training (tandem) data. Each tree contributes a vote as to the probability the given alignment is correct, and the final prediction is an average over the votes. This model is invariant under scaling transformations of features. Training is also efficient, which is important since models are tailored to the scenario at hand, and must be rebuilt anew each time Qtip runs. Finally, it is capable of reporting feature importances, which we examine in more detail in the context of our simulation experiments (Supplementary Note 12 and Supplementary Figures 4–9). Further details on the model are in Supplementary Note 13.

## Discussion

We presented the tandem simulation framework and the Qtip software tool implementing the framework. To date, strategies for predicting mapping qualities have either been *ad hoc* or have required the user to prepare training data tailored to the scenario at hand. Qtip runs alongside a read aligner and builds an input model, simulates tandem reads, aligns those using the same aligner and parameters, then uses the trained model to predict mapping qualities. The model and training data are produced automatically and are tailored to the scenario at hand. While Qtip adds overhead to the read alignment process, it reasonable, with time overhead in the 6–28% range and memory overhead in the 1– 10% range. This framework, its improved predictions and the evaluation performed here should make authors of downstream software tools more confident that mapping qualities can be treated as the probabilities they claim to be, and to integrate those probabilities into their models rather than simply thresholding.

Qtip’s predictions are accurate in various scenarios: various read lengths, unpaired or paired reads, various alignment tools & parameters, etc. We defined novel measures (RCA & RCE) and plots (CID & CSED) for evaluating and plotting mapping-quality predictions. The framework is easy to adapt to other aligners; the aligner must be modified to output feature data in an extra SAM field. Nor is it difficult to add new features to an already-adapted read aligner. Since Qtip’s ensemble tree model is scale-agnostic, scaling guesswork it not necessary when adding a feature.

This framework is also applicable to specialized alignment settings, such as spliced RNA-seq alignment. In that case, a nuanced notion of correctness is needed; we care not only where an alignment lands on the reference but also whether it includes the correct splice junctions. There is room for improvement in the area of predicting mapping qualities for spliced alignments. Popular tools use simplistic prediction functions drawing quality values from a small range of possibilities. TopHat [29] and STAR [30] report mapping quality of either 0 or 255 (repetitive versus unique) depending on the number of alignments found. Qtip’s approach would produce a full spectrum of values, potentially with large downstream benefits.

Tandem simulation works to the degree that tandem reads can be sampled “from the same distribution” as input reads. In reality, sampling from the same distribution is not possible. Qtip mimics some aspects of the input data but not others. Homopolymer extensions/retractions are not captured, for example, creating a fundamental difference between tandem and input reads. A tradeoff exists here: Qtip’s simple model mimics some aspects of the input without sacrificing efficiency, whereas a more complex and less efficient model could improve accuracy by mimicking more aspects. A task for future work is to measure various points in this tradeoff space, and to define measures for characterizing how and to what extent a set of tandem reads differs from the input reads.

A question for future work is whether Qtip’s sampling strategy can be improved. A strategy using importance sampling, for example, might favor tandem reads originating from more difficult-to-predict portions of the sample space. “Important” might originate from repetitive elements, or have certain patterns of mismatches and gaps. Together with appropriate weighting during model training, this could achieve comparable accuracy while reducing the number of tandem reads required. It could also reduce the prediction variability we see in experiments involving longer reads and more repetitive genomes.

## Acknowledgements

We thank Rafael Irizarry for helpful design discussions in the early stages of the project. We thank Abhinav Nellore and Michael Schatz for helpful comments on manuscript drafts. We thank Michael Schatz, Srividya Ramakrishnan, Maria Nattestad, Adam Phillippy and Sergey Koren for their help in obtaining and understanding the CHM1 Assemblytics results. BL was supported by NSF grant IIS-1349906 to BL, NIH/NIGMS grant R01GM118568 to BL, and NIH/NHGRI grant R01HG005220 to Rafael Irizarry.

## Availability of data and materials

The Qtip software is available in the GitHub repository https://github.com/BenLangmead/qtip. It is distributed under the open source MIT License. The version of the software evaluated in this manuscript is archived at DOI 0.5281/zenodo.556217.

The scripts and software used to perform the evaluation and draw the plots and tables in the manuscript are available in the GitHub repository https://github.com/BenLangmead/qtip-experiments. The version of the software used in this manuscript is archived at DOI 10.5281/zenodo.570957.

## Supplementary Note 1 Alignment heuristics in Bowtie 2

We elaborate our discussion of alignment heuristics with a few observations about how Bowtie 2 searches for alignments.

1. Only alignments at or above an *alignment score threshold S*_*min*_ are considered. These are called *valid*.
2. *Local similarity filters* require that one or more substrings of the read to align with few or no differences. For instance, a filter might require that alignments contain a length-*k* stretch of nucleotides that match the reference exactly. For long enough *k*, this disqualifies otherwise valid alignments.
3. *Repeat filters* avoid the extensive and mostly unproductive work performed when many candidates pass the local similarity filter. Say we take a 20 nt read then use a fast index lookup to find all reference locations where the first 10 nt of the read occur exactly. Say there are 100,000 such places. Instead of examining each candidate, Bowtie 2 will examine a subset and ignore the rest.
4. *Early stopping* halts alignment work once it seems to have become unproductive. For instance, the aligner might stop searching once a certain number of consecutive index queries have failed to yield a valid alignment.

## Supplementary Note 2 Prevalence of alignment errors

We defined three categories of alignment error:

1. The read is reported to have originated from a locus in the reference genome, but actually originates from a DNA sequence not included in the reference
2. No alignment to the reference is found, but the read actually originates from some locus in the reference
3. An alignment to locus *L*_*r*_ in the reference is reported, but the read actually originates from a different locus in the reference, *L*_*t*_

To measure prevalence of these errors in a realistic setting, we conducted three sets of simulation experiments. In all three, most simulated reads were derived from a known “target” reference genome, either human (GRCh38) or mouse (GRCm38), with genetic variation and sequencing errors added. These reads have their true point of origin in the reference genome. We also simulated reads from two types of foreign sequences, having their true point of origin outside the reference genome. The first type of foreign sequence consisted of “contaminating” species: species present in laboratories that have been shown in past studies to contaminate sequencing datasets. The second consisted of sequences from a human CHM1 hydatidiform mole assembly that were not present in the GRCh38 primary assembly, representing sequences present in a human donor but missing from the human reference, an important source of Category-1 error in practice [3]. All reads are aligned to the target reference only. The foreign sequences from the contaminating genomes are included in all experiments; the CHM1 sequences are included in one of the two human experiments, as indicated in Supplementary Table 1.

To obtain the first kind of foreign sequence, we simulated 7% of the reads from contaminating genomes: 1% each from mouse (GRCm38), the bacteria *Propionibacterium acnes*, *Mycolpasma Hominis*, *Mycolpasma fermentans*, *Acholeplasma laidlawii*, and *Mycolpasma hyorhinis*, and the fungus *Malassezia globosa*. We arrived at these contaminants after surveying studies on sequencing data contamination [5, 11, 13]. When a read from one of these genomes aligns to the GRCh38 primary assembly, we call this a Category-1a error.

For the two experiments that did not include foreign sequence from CHM1, we simulated the remaining 93% of reads from the reference genome. For the third experiment, we included CHM1-specific sequence by simulating 93% of the reads from a concatenation of the GRCh38 reference and a collection of sequences of length at least 260 present in the CHM1 assembly but not in GRCh38. Supplementary Note 3 gives further detail on how the CHM1-specific sequences were obtained. When a read from the CHM1 assembly aligns to GRCh38, we call this a Category-1b error.

We simulated four samples for each of the three experiments: unpaired and pairedend, 100-nt and 250-nt samples. We ran Mason as described in Supplementary Note 4. We aligned reads with Bowtie 2 v2.3.2 as described in Supplementary Note 5. In one experiment we inverted the roles of the mouse and human genomes, using mouse as the target and human as one of the contaminant genomes. For each aligned sample, we counted errors of the three categories and tabulated them as a percentage of the simulated reads and as a percentage of the errors (Supplementary Table 1).

As expected, longer reads yield fewer errors than shorter reads and paired-end reads yield fewer errors than unpaired reads. A large majority of the errors, ranging from 95.8-99.7%, are of category 3. The next largest, ranging from 0.2-3.9% of errors, are of category 2. These numbers could be reduced using more sensitive alignment parameters. Category 1 errors range from 0.27–0.46% of errors in the more realistic experiment that included CHM1 sequence, and from 0.05–0.014% in the other two experiments. The large fraction of category 3 errors underscores the need for better methods for predicting mapping quality.

**Supplementary Table 1:** Prevalence of errors of different categories when analyzing Mason-simulated samples using Bowtie 2. The “% reads” column gives percent of all the reads, both correctly and incorrectly aligned, having that error category, and “% errors” gives percent of errors having that error category.

| Target reference |  |  | Category 1a<br>(contamination) |  | Category 1b<br>(CHM1) |  | Category 2 |  | Category 3 |  |
| --- | --- | --- | --- | --- | --- | --- | --- | --- | --- | --- |
|  |  |  | % reads | % errors | % reads | % errors | % reads | % errors | % reads | % errors |
| GRCh38 | 100 | Unpaired | 0.006 | 0.144 | - | - | 0.159 | 3.567 | 4.287 | 96.289 |
|  |  | Paired | 0.004 | 0.136 | - | - | 0.079 | 2.657 | 2.888 | 97.206 |
|  | 250 | Unpaired | 0.002 | 0.100 | - | - | 0.065 | 2.713 | 2.325 | 97.187 |
|  |  | Paired | 0.002 | 0.094 | - | - | 0.073 | 3.902 | 1.794 | 96.004 |
| GRCh38+CHM1 | 100 | Unpaired | 0.006 | 0.144 | 0.009 | 0.205 | 0.164 | 3.692 | 4.266 | 95.960 |
|  |  | Paired | 0.004 | 0.136 | 0.005 | 0.178 | 0.078 | 2.622 | 2.884 | 97.064 |
|  | 250 | Unpaired | 0.002 | 0.099 | 0.009 | 0.363 | 0.067 | 2.758 | 2.348 | 96.780 |
|  |  | Paired | 0.002 | 0.093 | 0.003 | 0.180 | 0.073 | 3.884 | 1.809 | 95.843 |
| Mouse | 100 | Unpaired | 0.005 | 0.064 | - | - | 0.065 | 0.822 | 7.809 | 99.114 |
|  |  | Paired | 0.003 | 0.056 | - | - | 0.029 | 0.502 | 5.760 | 99.442 |
|  | 250 | Unpaired | 0.003 | 0.053 | - | - | 0.012 | 0.238 | 4.895 | 99.709 |
|  |  | Paired | 0.002 | 0.051 | - | - | 0.013 | 0.340 | 3.839 | 99.609 |

## Supplementary Note 3 Obtaining CHM1-specific sequences

We used MUMmer and nucmer [9] to align the MHAP CHM1 assembly [2] to the GRCh37 primary assembly. We analyzed the resulting alignment using Assemblytics [12], with results available at: http://assemblytics.com/analysis.php?code=human. We also downloaded the MHAP CHM1 assembly (accession: GCA 000772585) from Genbank [1].

Similarly, we used MUMmer and nucmer [9] to align the MHAP CHM1 assembly assembly to the GRCh38 primary assembly. We analyzed the resulting alignment using Assemblytics [12], with the results available at: http://assemblytics.com/analysis.php?code=sIOvqHUKNaCpU8NXdcTS. This assembly is like the one used above (accession: GCA 000772585), but with low-support contigs (having fewer than 50 supporting reads) removed. We also downloaded this version of the assembly from the University of Maryland at: http://gembox.cbcb.umd.edu/mhap/asm/human.quiver.ctg.fasta.gz.

We downloaded the Assemblytics results from both alignments by clicking the ”Download zip file of all results” link. Using the CHM1 assemblies and the structural variation BED file reported by Assemblytics, we extracted all insertions in the CHM1 assembly with respect to the reference genome with length at least 260. The minimum length restriction is required by Mason for simulations of 250 nt reads. We included 25 nt of overhang on either side of the extracted CHM1 sequences, to allow for reads aligning to the cusps. We constructed new “+CHM1” versions of the reference genomes by appending CHM1-specific sequences to the FASTA file containing the reference.

## Supplementary Note 4 Simulations

~~~
Mason. To generate unpaired samples we ran Mason v0.1.2 [7] with these parameters:
mason illumina -hn 2 -i -s (seed) -sq -n (read_len) -N 4000000 \
-o (output_fastq) (genome_fasta)
~~~

`illumina` causes Mason to simulate Illumina-like reads. `-hn 2` causes Mason to simulate 2 haplotypes. `-i` causes Mason to include information about the read’s true point of origin in the read name, which is useful later for determining which alignments are correct. `-s` sets the pseudo-random seed, which we set arbitrarily for our experiments. `-sq` causes Mason to simulate base qualities. `-n` specifies the read length. `-N` specifies the number of reads to simulate. `-o` specifies the output file.

To generate paired-end samples we ran Mason with these parameters:

~~~
mason illumina -hn 2 -i -s (seed) -sq -mp -rn 2 -ll (mean) -le (stddev) \
-n (read_len) -N 4000000 -o (output_fastq) (genome_fasta)
~~~

`illumina` causes Mason to simulate Illumina-like reads. `-hn 2` causes Mason to simulate 2 haplotypes. `-i` causes Mason to include information about the read’s true point of origin in the read name, which is useful later for determining which alignments are correct. `-s` sets the pseudo-random seed, which we set arbitrarily for our experiments. `-sq` causes Mason to simulate base qualities. `-mp` causes Mason to simulate paired-end reads. `-rn 2` enables “one-based slash suffix” read naming. `-ll` and `-le` are discussed below. `-n` specifies the read length. `-N` specifies the number of reads to simulate. `-o` specifies the output file.

For the paired-end samples, we ran Mason with parameters `-ll 3L -le L`. These establish a mean (3*L*) and standard deviation (*L*) for the fragment length distribution. This means that some simulated fragments will fall outside the range 2*L* − 4*L*, despite the fact that this is the range of fragment lengths that the aligners will consider “concordant” (Supplementary Note 5). This is desirable; for our paired-end samples, we would like the input to include reads that yield all three alignment categories: *conc*, *disc* and *bad-end*.

Mason v0.1.2 cannot handle very short reference sequences. To work around this, we removed all reference sequences shorter than 10,000 bases at the outset. We filtered out 90 short sequences from the GRCh38 assembly, totaling 229,926 nt. We filtered out one 4,262-nt-long sequence from the GRCh37 assembly. We filtered out one 1,976-nt-long sequence from the GRCm38 assembly. And we filtered out 5 short sequences from the AGPv4 assembly, totaling 37,762 nt.

When simulating paired-end reads, Mason v0.1.2 would sometimes generate pairs where the end that one would expect to find upstream would actually be found downstream. For example, in a typical Illumina paired-end protocols, the two ends of a fragment originating from the forward reference strand would align with (a) end 1 upstream of end 2, (b) end 1 aligning in the forward orientation, and (c) end 2 aligning in the reverse orientation. But for around 0.10–0.15% of the pairs in a Mason-simulated sample, we found that the upstream/downstream relationship between the ends was reversed. These were filtered out prior to our experiments.

**wgsim.** wgsim is one of the simulators we evaluate in Supplementary Note 8. To generate unpaired samples with wgsim, we ran it with these parameters:

~~~
wgsim -S (seed) -1 (read_len) -2 (read_len) -N 4000000 \ (genome_fasta) (output_fastq)
~~~

`-S` sets the pseudo-random seed, which we set arbitrarily for our experiments. `-1` and `-2` specify the read lengths for either end of a paired-end read. `-N` specifies the number of reads to simulate. The last two parameters specify the reference genome FASTA file and the output FASTQ file.

To generate paired-end samples we ran wgsim with these parameters:

~~~
wgsim -S (seed) -d (mean) -s (stddev) -1 (read_len) -2 (read_len) \
-N 4000000 (genome_fasta) (output_fastq_1) (output_fastq_2)
~~~

`-S` sets the pseudo-random seed, which we set arbitrarily for our experiments. `-d` sets the mean fragment length and `-s` sets the fragment length standard deviation. `-1` and `-2` specify the read lengths for either end of a paired-end read. `-N` specifies the number of reads to simulate. The last two parameters specify the reference genome FASTA file and the output FASTQ files.

**Art.** Art is one of the simulators we evaluate in Supplementary Note 8. To generate unpaired samples with Art, we ran it with these parameters:

~~~
art_illumina -sam -na -rs (seed) -l (read_len) -f (fold_coverage) \
-i (genome_fasta) -o (output_fastq)
~~~

`-sam` enables SAM output. Information about a simulated read’s point of origin is encoded in the SAM file. `-na` suppresses the ALN alignment file. `-rs` sets the pseudorandom seed, which we set arbitrarily for our experiments. `-l` specifies the read length. `-f` specifies the fold coverage. We set this to be high enough that we are eventually able to obtain 4 million reads or pairs per sample. `-i` specifies the reference genome FASTA file. `-o` specifies the output file.

To generate paired-end samples we ran Art with these parameters:

~~~
art_illumina -sam -na -p -rs (seed) -m (mean) -s (stddev) -l (read_len) \
-f (fold_coverage) -i (genome_fasta) -o (output_fastq)
~~~

`-sam` enables SAM output. Information about a simulated read’s point of origin is encoded in the SAM file. `-na` suppresses the ALN alignment file. `-rs` sets the pseudorandom seed, which we set arbitrarily for our experiments. `-m` specifies the mean fragment length. `-s` specifies the standard deviation. `-l` specifies the read length. `-f` specifies the fold coverage. We set this to be high enough that we are eventually able to obtain 4 million reads or pairs per sample. `-i` specifies the reference genome FASTA file. `-o` specifies the output file.

## Supplementary Note 5 Read alignment

### Bowtie 2

For paired-end samples, we ran Bowtie 2 with the `-I` and `-X` parameters to set the minimum and maximum concordant fragment lengths. We specified `-I 2L -X 4L` where `L` corresponds to the `-ll` and `-le` parameters specified when running Mason. For experiments where we varied sensitivity level, we also specified a sensitivity parameter: `--very-fast`, `--fast`, `--sensitive` or `--very-sensitive`. For experiments where Bowtie 2 was run in local alignment mode, we also specified the `--local` parameter. Additionally, Qtip always specifies the `--mapq-extra` option when running Bowtie 2, which enables feature-reporting features. When running Qtip together with Bowtie 2, the user may also specify additional Bowtie 2 parameters on the command line.

### BWA-MEM

For paired-end samples, we ran BWA-MEM with the `-I` option to set the minimum and maximum concordant fragment lengths. BWA-MEM’s default behavior, unlike either Bowtie 2 or SNAP, is to infer the fragment length distribution over a sample of the input reads. By specifying `-I` we disabled that automatic inference. This was to ensure that the read aligners worked similarly and, in particular, that all agreed on which fragment lengths should be considered concordant. The `-I` parameter sets the mean and standard deviation of the fragment length distribution. BWA-MEM consider any fragment length within 4 standard deviations of the mean to be concordant. Thus, we set `-I 3L,L/4`, achieving the same result as we did above with Bowtie 2’s `-I` and `-X` parameters. When running Qtip together with BWA-MEM, the user may also specify additional BWA-MEM parameters on the command line.

### SNAP

For paired-end samples, we ran SNAP with the `-s` parameter to set the minimum and maximum concordant fragment lengths. We specified `-s L 3L` indicating that the distance between the leftmost base of the upstream made and the leftmost base of the downstream made is allowed to vary from *L* to 3*L*, achieving the same result as we did above with Bowtie 2’s `-I` and `-X` parameters. Additionally, Qtip always sets SNAP’s `-=` parameter for reasons described in Supplementary Note 11. When running Qtip together with SNAP, the user may also specify additional SNAP parameters on the command line.

## Supplementary Note 6 Modifications to read aligners

We modified three alignment tools — Bowtie 2, BWA-MEM, and SNAP— to output feature data needed by Qtip. To make the modifications, we studied each tool’s source code and identified data that could be relevant to the mapping quality prediction task. We started by studying how the tool itself predicts mapping quality, since the inputs to those routines are candidate features. In all three cases, we added more features beyond these, usually with the goal of capturing more information about the repetitiveness of the read and how many of its “seeds” either failed to align or aligned repetitively.

Here we examine each tool, discussing (a) how the tool predicts mapping quality, (b) which features we re-used from that process, and (c) which additional features we added.

### Bowtie 2

Bowtie 2’s mapping quality calculation uses a lookup table, with the lookup keys being functions of (a) the best alignment’s score, (b) the second-best alignment’s score, (c) whether end-to-end or local alignment were used, (d) whether the read aligned concordantly in a pair. We preserve (a) and (b) in the features we elect to report from Bowtie 2. We do not need to report either (c) or (d) because these follow naturally from how Qtip works. In the case of (c), the training data is tailored to match the alignment parameters used to align the input data; the training data will always match the input data in terms of the alignment mode used. In the case of (d), Qtip trains separate models for separate alignment categories (*conc*, *disc*, *unp*, *bad-end*), and always uses the appropriate model for prediction.

Following is a list of the unpaired features output in the modified version of Bowtie 2:

- Best alignment score
- Difference between the best and second-best alignment score, or NA if no second-best alignment was found
- Difference between the best and second-best edit distance, or NA if no second-best alignment was found
- Number of seeds for which Bowtie 2 sought an alignment (*attempted* seeds) divided by the number of nucleotides in the read
- Fraction of attempted seeds that aligned uniquely (to exactly one place in the reference)
- Same as previous but limited to seeds that aligned in the same orientation as the overall reported alignment
- Fraction of attempted seeds that aligned repetitively (to more than one place in the reference)
- Same as previous but limited to seeds that aligned in the same orientation as the overall reported alignment
- Mean number of alignments found for any attempted seed
- Same as previous but limited to seeds that aligned in the same orientation as the overall reported alignment

Following is a list of the paired-end features output in the modified version of Bowtie 2:

- Sum of the alignment scores for both ends of the best paired-end alignment
- Difference between the best and second-best paired-end alignment score (i.e. the sum of the scores of the two ends), or NA if no second-best alignment was found
- Difference between the best and second-best paired-end edit distance (i.e. the sum of the edit distances of the two ends), or NA if no second-best alignment was found

Supplementary Note 12 discusses the relative importance of these features as we measured in our simulation experiments.

### BWA-MEM

BWA-MEM’s mapping quality calculation depends on, among other factors, (a) the best alignment’s score, (b) the second-best alignment’s score, (c) the number of alignments “tied” for second-best, (d) the number of aligned bases (i.e. excluding any soft-clipped bases), (e) the percent identity, summarized across the aligned bases, (f) the *seed coverage*, which roughly equals the mean number of times a read base is covered by an aligned seed. We use these features and others in Qtip.

Following is a list of the unpaired features output in the modified version of BWA-MEM :

- Best alignment score
- Edit distance of the best alignment
- Difference between the best and second-best alignment score, or NA if no second-best alignment was found
- Number of alignments “tied” for second-best
- Fraction of seeds that aligned repetitively
- Seed coverage

Following are the paired-end features output in the modified version of BWA-MEM :

- Difference between the best and second-best paired-end alignment score (i.e. the sum of the scores of the two ends), or NA if no second-best alignment was found
- Sum of alignment scores of the two ends of the best paired-end alignment
- Number of paired-end alignments “tied” for second-best
- Fraction of seeds from either end of the pair that aligned repetitively
- Seed coverage over both ends of the pair

Supplementary Note 12 discusses the relative importance of these features as we measured in our simulation experiments.

### SNAP

For a given read, SNAP assigns a probability to each alignment found for that read. SNAP’s procedure for calculating the best alignment’s mapping quality mostly consists of dividing the best alignment’s probability by the sum of the probabilities of all the other alignments found. The probability of a given alignment is function of the length of the read, the read’s quality values, and a collection of parameters giving the prior probability of a variant, gap opening, gap extension, etc. We use these probabilities and other features in Qtip.

Following are the unpaired features we output in the modified version of SNAP:

- Edit distance of the best alignment
- Difference between best and second-best alignment’s edit distance, or NA if no second-best alignment was found
- Probability of the best alignment divided by the total probability of all the alignments found
- Number of seeds ignored because they matched in the genome too many times
- Minimum number of times a matching seed matched in the genome
- Number of seeds that failed to match the genome
- Mean number of matches per seed

Following are the paired-end features we output in the modified version of SNAP:

- Difference between the best and second-best paired-end edit distance (i.e. the sum of the scores of the two ends), or NA if no second-best alignment was found
- Total number of seeds in the pair ignored because they matched in the genome too many times
- Minimum number of times a matching seed matched in the genome across both ends of the pair
- Number of seeds that failed to match the genome across both ends of the pair
- Mean number of matches per seed across all seeds in the pair

Supplementary Note 12 discusses the relative importance of these features as we measured in our simulation experiments.

Qtip parses feature data from the extra `ZT:Z` SAM field. So once we identified a feature, we enable Qtip to use that feature simply by printing it as an additional value in the `ZT:Z` field. This field contains comma-separated numeric data. The following is an example of an alignment produced by BWA-MEM with the `ZT:Z` field. The `ZT:Z` field is not printed for unaligned reads, and Qtip will ignore these.

Qtip also ignores the small number of chimeric alignments reported by BWA-MEM. Proper handling of chimeric alignments is future work.

In addition to the features discussed above, which are reported by the read aligners in the `ZT:Z` field, Qtip adds a few additional features, specifically:

- Length of the read or end
- Sum of the base qualities of the aligned bases in the read or end
- Sum of the base qualities of the soft-clipped bases in the read or end

Finally, for a given end of a paired-end alignment, whether concordant or discordant, feature data for the opposite end is included in the record for the given end. I.e., in the record for a given end of a paired-end read, you will find aligner-specific unpaired feature data, aligner-specific paired-end feature data, generic features, and both aligner-specific and generic features for the opposite end.

When the training and test data are compiled within Qtip, columns in which all elements are equal are eliminated. E.g., if the input consists of reads all of the same length, then the read length will be identical for all records and the corresponding column will be dropped from the training and test matrices. Similarly, if the input consists of reads aligned in end-to-end alignment mode (Bowtie 2’s default), then the sum of the base qualities of the soft-clipped bases will always be 0, and the column corresponding to that feature will again be dropped.

### Availability of aligner modifications

Bowtie 2 v2.3.2 has the appropriate feature-printing code built in; earlier versions of Bowtie 2 do not have this code. The `--mapq-extra` parameter instructs Bowtie 2 v2.3.2 to print feature data in the `ZT:Z` extra SAM field. To run this version of Bowtie 2 in combination with Qtip, use Qtip’s `--bt2-exe` parameter.

Qtip comes with a source-code patch that causes BWA-MEM to output feature data for use with Qtip. To create a modified version of BWA-MEM, download the BWA-MEM v0.7.15 source archive from https://sourceforge.net/projects/bio-bwa/files/. Once extracted, change to the BWA-MEM v0.7.15 source directory and run:

~~~
patch -p1 < /path/to/qtip/software/bwa/bwa_conc_flags_0.7.15.patch
~~~

Then build the binaries as usual. To run this version of BWA-MEM in combination with Qtip, use Qtip’s `--bwa-exe` parameter. The modified BWA-MEM will print the feature data in the `ZT:Z` extra SAM field.

Qtip comes with a source-code patch that causes SNAP to output feature data for use with Qtip. To create a modified version of SNAP, download the SNAP v1.0beta.18 source archive from http://snap.cs.berkeley.edu. Once extracted, change to the SNAP v1.0beta.18 source directory and run:

~~~
patch -p1 < /path/to/qtip/software/snap/snap_features_patch.diff
~~~

Then build the binaries as usual. To run this version of SNAP in combination with Qtip, use Qtip’s `--snap-exe` parameter. The modified SNAP will print the feature data in the `ZT:Z` extra SAM field.

**Supplementary Table 2:** Relative change in area under CID (RCA) and relative change in sum of squared error (RCE) for various aligners and two versions of the human reference genome, expressed as percent change. Some experiments simulated reads from just the reference (“GRCh37,” “GRCh38”) while other simulated from insertions in the CHM1 assembly relative to the reference assembly (“GRCh37+CHM1,” “GRCh38+CHM1”). The experiments used 100 nt or 250 nt reads, and unpaired or paired-end reads, as indicated. Results are means and standard deviations over 10 random trials, repeated starting from the input modeling step.

|  |  |  | 100 nt |  |  |  | 250 nt |  |  |  |
| --- | --- | --- | --- | --- | --- | --- | --- | --- | --- | --- |
|  |  |  | RCA |  | RCE |  | RCA |  | RCE |  |
|  |  |  | mean | sd | mean | sd | mean | sd | mean | sd |
| Unpaired | Bowtie 2 | GRCh37 | -11.22 | 1.08 | -24.43 | 0.66 | -15.02 | 0.39 | -28.37 | 0.38 |
|  |  | GRCh37+CHM1 | -10.77 | 0.71 | -24.22 | 0.35 | -12.78 | 2.07 | -27.67 | 1.14 |
|  |  | GRCh38 | -7.43 | 1.82 | -18.53 | 2.61 | -16.82 | 2.01 | -19.97 | 0.44 |
|  |  | GRCh38+CHM1 | -7.39 | 1.31 | -18.50 | 1.26 | -16.36 | 2.03 | -19.76 | 1.07 |
|  | BWA-MEM | GRCh37 | -14.49 | 2.29 | -49.54 | 0.43 | -9.14 | 2.09 | -52.31 | 0.38 |
|  |  | GRCh37+CHM1 | -15.55 | 0.81 | -49.34 | 0.29 | -11.62 | 6.27 | -51.42 | 0.35 |
|  |  | GRCh38 | -15.49 | 0.58 | -47.42 | 0.37 | -15.78 | 0.57 | -51.14 | 0.31 |
|  |  | GRCh38+CHM1 | -15.53 | 0.56 | -47.14 | 0.33 | -15.57 | 0.79 | -50.80 | 0.54 |
|  | SNAP | GRCh37 | -15.94 | 0.32 | -36.86 | 0.23 | -9.53 | 3.88 | -28.57 | 0.47 |
|  |  | GRCh37+CHM1 | -16.39 | 0.93 | -36.95 | 0.29 | -6.07 | 6.84 | -27.66 | 0.44 |
|  |  | GRCh38 | -19.58 | 0.18 | -36.58 | 0.47 | -14.74 | 0.27 | -25.47 | 0.32 |
|  |  | GRCh38+CHM1 | -19.51 | 0.24 | -36.69 | 0.37 | -14.75 | 0.32 | -25.29 | 0.71 |
| Paired | Bowtie 2 | GRCh37 | -48.62 | 0.23 | -65.07 | 0.25 | -54.09 | 1.55 | -68.39 | 0.31 |
|  |  | GRCh37+CHM1 | -46.72 | 0.59 | -64.70 | 0.14 | -53.18 | 1.36 | -68.06 | 0.32 |
|  |  | GRCh38 | -34.93 | 1.00 | -54.32 | 0.19 | -47.93 | 0.26 | -57.77 | 0.19 |
|  |  | GRCh38+CHM1 | -34.92 | 0.93 | -54.28 | 0.21 | -47.07 | 0.35 | -57.54 | 0.19 |
|  | BWA-MEM | GRCh37 | -13.33 | 0.23 | -45.70 | 0.27 | -10.08 | 0.75 | -47.58 | 0.31 |
|  |  | GRCh37+CHM1 | -14.21 | 0.70 | -45.02 | 0.39 | -15.31 | 1.41 | -46.90 | 0.36 |
|  |  | GRCh38 | -14.19 | 0.16 | -41.35 | 0.19 | -12.36 | 0.46 | -42.78 | 0.29 |
|  |  | GRCh38+CHM1 | -13.89 | 0.23 | -40.97 | 0.25 | -12.68 | 0.33 | -42.52 | 0.27 |
|  | SNAP | GRCh37 | -56.53 | 1.99 | 1.39 | 2.34 | -42.89 | 7.63 | 13.17 | 3.95 |
|  |  | GRCh37+CHM1 | -54.39 | 1.67 | 1.79 | 1.73 | -42.30 | 8.09 | 14.39 | 5.07 |
|  |  | GRCh38 | -51.36 | 0.98 | -11.16 | 1.12 | -51.32 | 1.45 | 4.34 | 2.29 |
|  |  | GRCh38+CHM1 | -51.11 | 0.97 | -11.43 | 0.86 | -50.91 | 1.48 | 3.88 | 2.06 |

## Supplementary Note 7 Varying aligner sensitivity

We used Qtip together with Bowtie 2 to align and predict mapping qualities for the Mason-simulated 100 nt and 250 nt unpaired and paired-end samples. We repeated each experiment four times, varying Bowtie 2’s sensitivity parameter: `--very-fast`, `--fast`, `--sensitive` or `--very-sensitive`. We also tried Qtip both in its (default) end-to-end alignment mode and in its local alignment mode. We calculated RCA and RCE for all 10 trials of all experiments (Supplementary Table 3) and plotted CSED for the first trial of the experiments that used the 100 nt reads (Supplementary Figure 1).

Qtip-predicted mapping qualities are superior overall to Bowtie 2’s in every scenario tested, as indicated by the negative RCAs and RCEs (Supplementary Table 3). Variability of RCAs and RCEs across trials is generally modest, but higher for `--very-sensitive` mode. Standard deviations range up to 5.03.

CSED curves (Supplementary Figure 1) again show that for some ranges of cutoffs, aligner-reported mapping qualities are better at minimizing cumulative squared error, corresponding to the portions of the CID curves that rise above *y* = 0. This is most clearly observed for unpaired `--very-sensitive` alignment. But Qtip provides superior predictions for the vast majority of cutoffs, including nearly all cutoffs for the paired-end experiments.

**Supplementary Table 3:** Relative change in area under CID (RCA) and relative change in sum of squared error (RCE) when running Qtip and Bowtie 2 at various levels of sensitivity on Mason-simulated Illumina-like samples. Relative change is expressed as a percent. Each simulated sample consists of 4 million reads or pairs. Bowtie 2 is run in either endto-end or local alignment mode as indicated. Results are means and standard deviations over 10 random trials, repeated starting from the input modeling step.

|  | Length | Sensitivity | End-to-end |  |  |  | Local |  |  |  |
| --- | --- | --- | --- | --- | --- | --- | --- | --- | --- | --- |
|  |  |  | RCA |  | RCE |  | RCA |  | RCE |  |
|  |  |  | mean | sd | mean | sd | mean | sd | mean | sd |
| Unpaired | 100 | –very-fast | -9.55 | 1.04 | -20.73 | 1.13 | -17.06 | 0.37 | -47.04 | 0.23 |
|  |  | –fast | -8.33 | 0.28 | -19.96 | 0.26 | -13.66 | 0.37 | -36.52 | 0.42 |
|  |  | –sensitive | -7.51 | 1.47 | -18.74 | 1.57 | -11.58 | 0.66 | -29.62 | 0.45 |
|  |  | –very-sensitive | -6.53 | 1.65 | -18.32 | 2.62 | -9.01 | 1.78 | -21.75 | 4.61 |
|  | 250 | –very-fast | -25.26 | 0.77 | -32.32 | 0.43 | -32.44 | 1.59 | -52.37 | 0.40 |
|  |  | –fast | -18.75 | 1.63 | -24.21 | 0.71 | -24.16 | 1.40 | -35.51 | 0.60 |
|  |  | –sensitive | -16.38 | 2.07 | -19.25 | 1.80 | -22.27 | 3.55 | -29.07 | 1.12 |
|  |  | –very-sensitive | -16.10 | 2.64 | -17.58 | 2.40 | -15.59 | 5.03 | -20.87 | 1.89 |
| Paired | 100 | –very-fast | -32.56 | 1.00 | -50.48 | 0.26 | -67.65 | 0.21 | -49.77 | 0.63 |
|  |  | –fast | -33.69 | 0.51 | -51.75 | 0.22 | -66.36 | 0.12 | -51.48 | 0.51 |
|  |  | –sensitive | -35.56 | 0.65 | -54.34 | 0.12 | -63.63 | 0.33 | -52.06 | 0.36 |
|  |  | –very-sensitive | -37.00 | 0.32 | -56.43 | 0.09 | -60.54 | 0.30 | -53.97 | 0.63 |
|  | 250 | –very-fast | -46.44 | 0.45 | -57.22 | 0.25 | -71.51 | 0.60 | -62.20 | 1.86 |
|  |  | –fast | -45.04 | 3.03 | -58.18 | 0.19 | -69.35 | 0.34 | -63.67 | 1.36 |
|  |  | –sensitive | -47.42 | 0.72 | -57.80 | 0.14 | -67.00 | 1.35 | -61.53 | 2.56 |
|  |  | –very-sensitive | -46.76 | 1.70 | -57.93 | 0.22 | -63.41 | 0.55 | -62.27 | 1.48 |

**Supplementary Figure 1:**
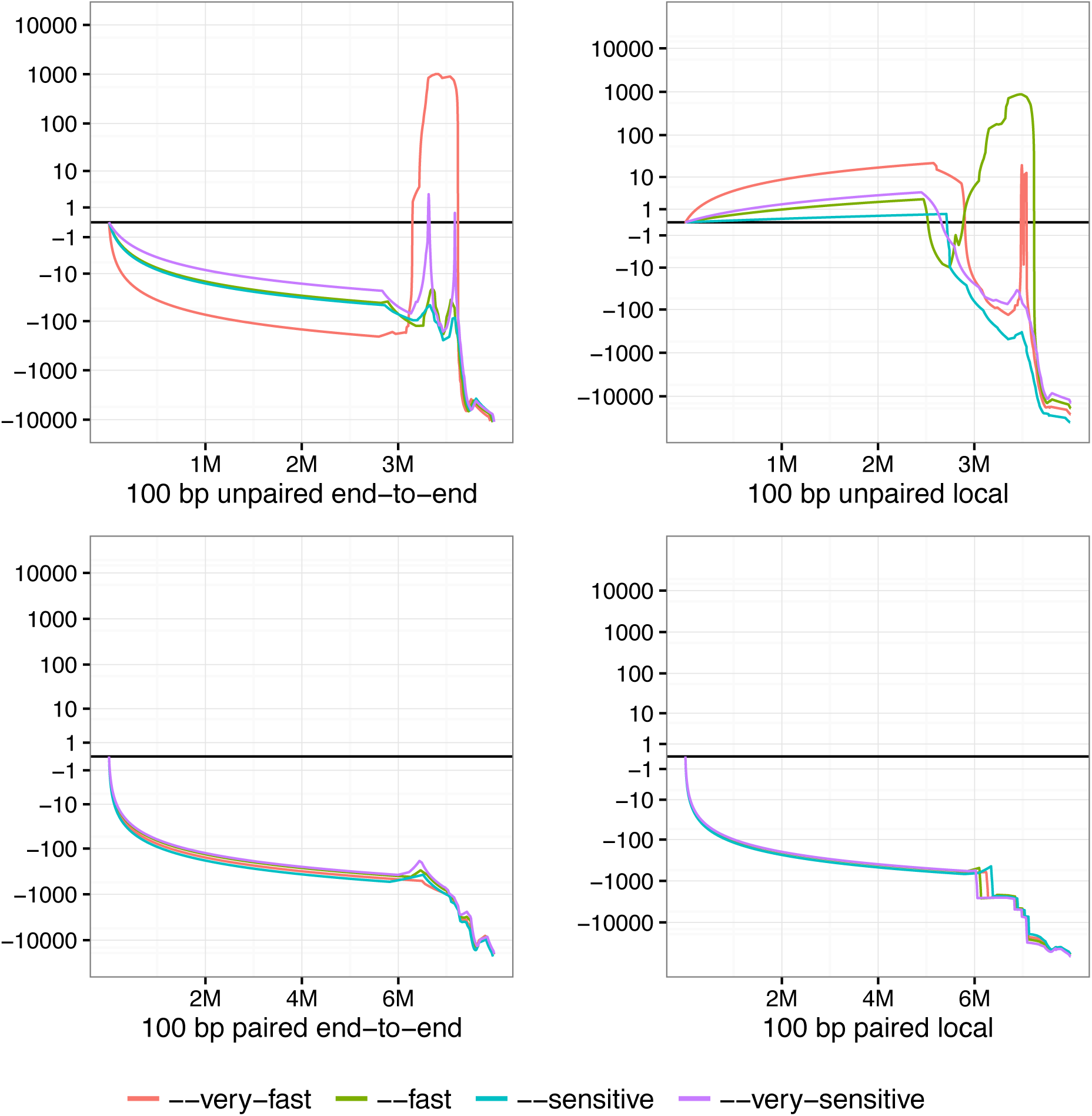
Cumulative squared-error difference (CSED) plot from running Qtip and Bowtie 2 and varying Bowtie 2’s sensitivity level. The input reads are Mason-simulated Illumina-like 100 nt samples, each consisting of 4 million reads/pairs. The horizontal axis indicates cumulative number of reads/ends passing the cutoff, with the left-hand extreme corresponding to a high mapping-quality cutoff and right-hand extreme corresponding to a low cutoff. Results for unpaired samples are shown on top, and paired on bottom. Bowtie 2 is run in its (default) end-to-end mode in the case of the left-hand plots, and in local mode in the case of the right-hand plots.

## Supplementary Note 8 Varying read simulator

Different read simulators make different decisions about how to emulate the sequencing process and its errors and biases. To study how this impacts Qtip’s predictions, we ran a series of experiments using Mason [7] and the popular wgsim (https://github.com/lh3/wgsim) and Art [8] simulators. For all three, we simulated 100 nt and 250 nt samples, both unpaired and paired-end. See Supplementary Note 4 for the commands used and details about simulated fragment lengths. We then used Qtip together with Bowtie 2 to align and predict mapping qualities. We calculated RCA and RCE for all 10 trials of all experiments (Supplementary Table 4) and plotted CID for the first trial (Supplementary Figure 2). Overall, the results show the choice of simulator does not have a large effect on the outcome of the experiment. Differences in prediction performance on the paired-end samples may be due in part to the different fragment length distributions generated by the simulators, as discussed in Supplementary Note 4.

**Supplementary Figure 2:**
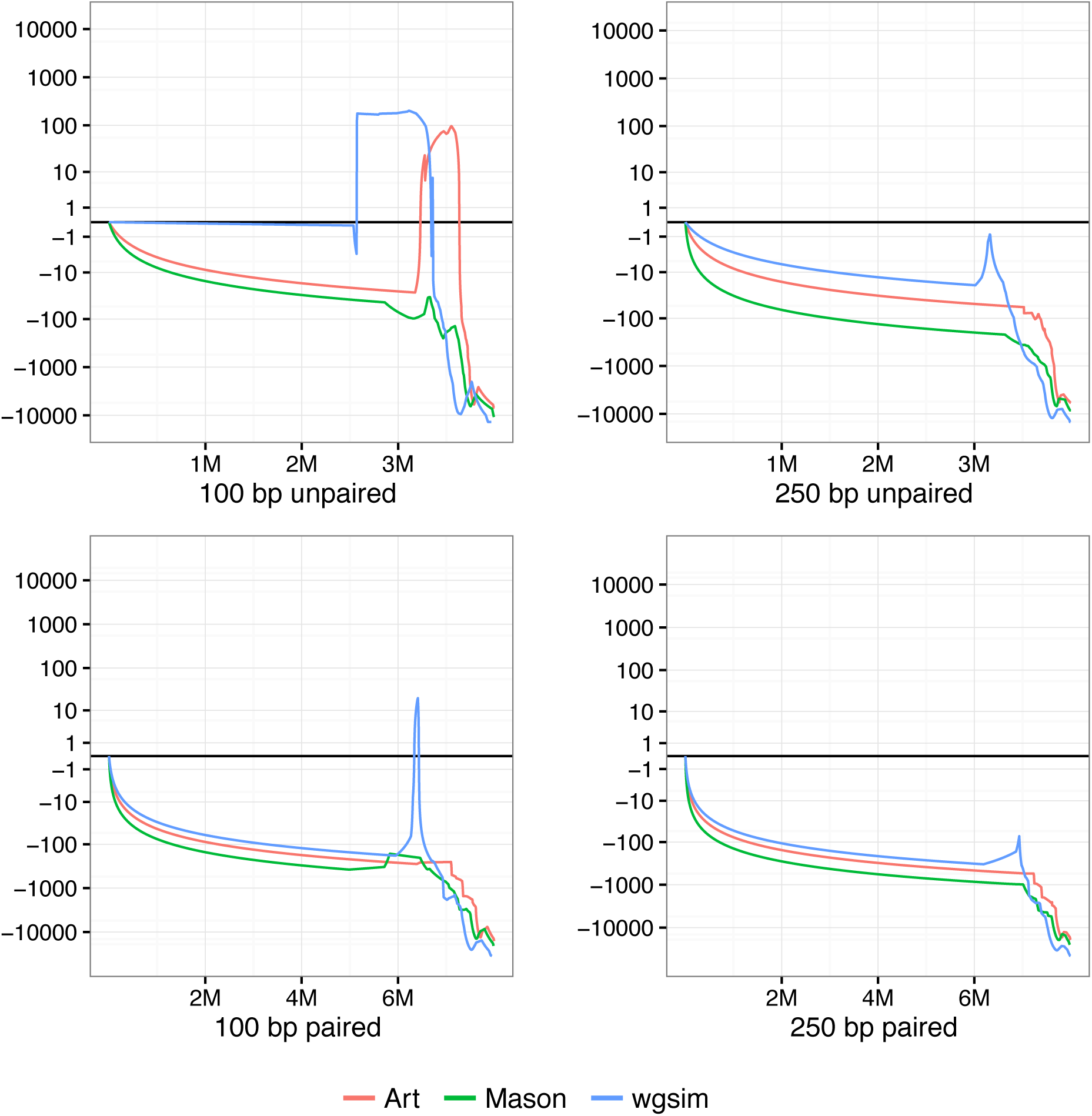
Cumulative squared-error difference (CSED) plot from running Qtip and Bowtie 2 on samples simulated from three different read simulators: Mason, wgsim and Art. Each sample consists of 4 million reads or pairs. The horizontal axis indicates cumulative number of reads/ends passing the cutoff, with the left-hand extreme corresponding to a high quality cutoff and right-hand extreme to a low cutoff.

**Supplementary Table 4:** Relative change in area under CID (RCA) and relative change in sum of squared error (RCE) for samples simulated by various simulators. Bowtie 2 endto-end alignment was used in all cases. Relative change is expressed as a percent. The experiments used 100 nt or 250 nt reads, and unpaired or paired-end reads, as indicated. Results are means and standard deviations over 10 random trials, repeated starting from the input modeling step.

|  |  |  | RCA |  | RCE |  |
| --- | --- | --- | --- | --- | --- | --- |
|  |  |  | mean | sd | mean | sd |
| Unpaired | 100 nt | Art | -6.26 | 0.35 | -15.87 | 0.66 |
|  |  | Mason | -7.43 | 1.82 | -18.53 | 2.61 |
|  |  | wgsim | -9.06 | 0.25 | -19.60 | 0.90 |
|  | 250 nt | Art | -12.41 | 0.84 | -19.83 | 0.53 |
|  |  | Mason | -16.82 | 2.01 | -19.97 | 0.44 |
|  |  | wgsim | -18.52 | 0.32 | -26.93 | 0.50 |
| Paired | 100 nt | Art | -26.02 | 0.83 | -41.98 | 0.26 |
|  |  | Mason | -34.93 | 1.00 | -54.32 | 0.19 |
|  |  | wgsim | -26.37 | 0.84 | -46.00 | 0.19 |
|  | 250 nt | Art | -40.37 | 0.36 | -47.21 | 0.30 |
|  |  | Mason | -47.93 | 0.26 | -57.77 | 0.19 |
|  |  | wgsim | -39.18 | 1.10 | -54.41 | 0.21 |

## Supplementary Note 9 Variant calling experiment

### Alignment

We used Qtip v1.6.2 together with Bowtie 2 v2.3.2 to align the paired-end sequencing reads from ERR194147. The Qtip command used was:

~~~
qtip \
--ref ${FASTA_PATH}/hg38.fa \
--m1 ERR194147_1.fastq --m2 ERR194147_2.fastq \
--index ${INDEX_PATH}/hg38.fa \
--bt2-exe ${BOWTIE2_PATH}/bowtie2 \
--keep-intermediates \
--output-directory ${ODIR} \
--write-orig-mapq \
--write-precise-mapq \
--temp-directory ${TEMP} \
-${BOWTIE2_ARGS}
~~~

The Bowtie 2 arguments used (inserted where `${BOWTIE2_ARGS}` appears above) are: `-I 0 -X 550 -t -p 24`. The first two parameters set minimum and maximum fragment length to 0 and 550 respectively. The `-t` parameter enables extra timing output. The `-p 24 --reorder` parameters run Bowtie 2 with 24 threads while keeping output alignments in an order corresponding to the input reads.

With these parameters, the output directory from Qtip contains both an `input.sam` file with the original mapping qualities, and a `final.sam` file with the Qtip-predicted mapping qualities for the same alignments.

### Variant calling

After converting the SAM output from the alignment run to sorted BAM using sambamba [15], we then used Freebayes v1.1.0 to call variants:

~~~
freebayes \
-X -u --haplotype-length 0 \
-f ${FASTA_PATH}/hg38.fa \
--min-mapping-quality ${MIN_MAPQ} \
-v ${OUTPUT_VCF} \
${INPUT_BAM}
~~~

The `-X -u` arguments ensure Freebayes calls only SNVs and indels, though the indels calls were not studied here. The `--haplotype-length 0` argument ensures each SNV appears as a separate call in the VCF file, even in cases where nearby SNVs can be phased into a haplotype block. This helps make the SNVs more comparable between the Freebayes output and the Platinum calls. `-f` specifies the reference FASTA file. `--min--mapping-quality` specifies the minimum mapping quality an alignment must have to be included in the variant calling analysis. We run Freebayes repeatedly setting different values for `${MIN_MAPQ}`. We also tried running Freebayes in three other modes affecting the mapping quality threshold: (a) its default mode, (b) `--standard-filters` mode, and (c) `--use-mapping-quality`. But for none of the *β*s tested did the cutoffs enforced by any of those three modes yield maximal *F*_*β*_. In fact, the thresholds yielding maximal *F*_*β*_ were always in the range 2–5.

The above command was run twice for every mapping quality threshold: once with `${INPUT_BAM}` is set to the sorted BAM derived from the `input.sam` file, and once with with `${INPUT_BAM}` set to the sorted BAM derived from `input.sam`.

### Variant filtering

We applied two filters to the variant calls output by Freebayes. First, we eliminated variant calls with spuriously high coverage, as suggested in prior work [10]. Spuriously high coverage tends to indicate the presence of a copy number change that confounds variant calling. This was accomplished with a simple `awk` script that removed VCF records with depth (as indicated by the `DP` field of the `INFO` column) at least four poisson standard deviations above the mean. Mean depth of coverage was calculated to be 52.79 using `samtools flatstat`, so the filter removed variants with depth greater than 82. This script, and all others used for these experiments, can be found at https://github.com/BenLangmead/qtip-experiments/tree/v1.6.1.

Next, we filtered the Freebayes output to eliminate variant calls outside genomics regions deemed “high confidence” by the Platinum Genomes [4] project. We used the `vcfintersect` tool from the `vcflib` (https://github.com/vcflib/vcflib) collection; the specific command was:

~~~
vcfintersect -b ${PLATINUM_BED} ${VCF}
~~~

Where `${VCF}` was the file produced by Freebayes and `${PLATINUM_BED}` was a version of the Platinum Genomes high-confidence region `bed` file obtained from the Platinum Genomes FTP site (ftp://ussd-ftp.illumina.com/2016-1.0/hg38/small_variants/). We first preprocessed the `${PLATINUM_BED}` file to retain only regions in chromosomes 1–22 and X.

**F scores:** To obtain F-scores we compiled two `roc` files, one for the variant calls using the aligner-predicted mapping qualities and one for the calls predicted by Qtip. A `roc` file summarizes the number of true positives, false positives, and false negatives at all possible `QUAL` cutoffs. The `QUAL` field of the VCF file gives Freebayes’ “genotype confidence.” The ROCs were obtained by running `vcfroc`, a tool in the `vcflib` (https://github.com/vcflib/vcflib) collection. The `vcfroc` command was:

~~~
vcfroc \
--truth-vcf ${PLATINUM_VCF} \
--reference ${FASTA_PATH}/hg38.fa \
${VCF_FN} > ${ROC_FN}
~~~

### Finding the maximal F scores

All of the above experiments were carried out for all mapping-quality thresholds and for both the original and the Qtip-predicted mapping qualities. For the original mapping qualities and given a particular *β*, we calculated maximal *F*_*β*_ by evaluating *F*_*β*_ :

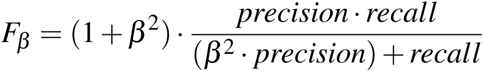

For every possible combination of mapping-quality and genotype-quality threshold. We then reported the maximal *F*_*β*_.

## Supplementary Note 10 Setting the number of tandem reads

Qtip has parameters that control the minimum number of tandem reads or pairs of each category (*conc*, *disc*, *bad-end* and *unp*) to generate. The default number of training reads/pairs generated for each category is 45. 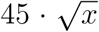 where *x* is the number of input alignments of that category. Both the scaling factor and the function are configurable via Qtip’s `--sim-function` and `--sim-factor` parameters.

We conducted a set of simulation experiments on a few selected combinations of functions and coefficients. We used Mason to simulate two sets of 50M reads, one unpaired set and one paried-end. Both sets used 100 nt reads, with paired-end reads having simulated fragments lengths mostly between 200 and 400 nt, per the strategy described in Supplementary Note 4. We tried the following policies for setting a target number of tandem alignments:

1. 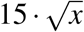
2. 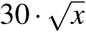
3. 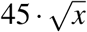
4. 0.01 · *x*
5. 0.03 · *x*
6. 0.05 · *x*
7. 50, 000
8. 100, 000

We say that these policies set a “target” number of alignments because the actual tandem simulation procedure could result in fewer or more alignments. This is because (a) not every tandem read will align successfully, (b) the target is inflated somewhat prior to simulation in order to account for alignment failures, and (c) the tandem simulator actually conducts a large number of binomial draws; the total across all the draws will be very close to, but not necessarily equal to, the inflated target.

We tried each of these policies for each of the following numbers of input reads:

1. 1,000,000
2. 5,000,000
3. 10,000,000
4. 50,000,000

The script used to drive these experiments is present in the `qtip-experiments` repository as experiments/simulated reads/train series.sh.

As seen in Supplementary Figure 3, the formulae result in similar levels accuracy as the number of input reads grows to 50,000,000, with an RCE spread of less than 2 in the unpaired case and less than 1 in the paired-end case. Setting the target to 50, 000 performs worst at at the highest numbers of input reads, suggesting the constant functions tend to perform poorly as the number of input alignment grows large. The linear functions (reddish lines) and those that grow as a function of the square root of the number of input alignments (greenish lines) perform similarly, though the 0.01 · *x* target performs poorly with only 1,000,000 input reads.

Ultimately, we prefer the formulae that grow with the square root of the number of input reads, as they represent a compromise between constant functions that perform poorly at higher numbers and linear functions that require large amounts of training data at larger input sizes. The square-root functions yield comparable RCE to the linear functions. Qtip’s default formula is 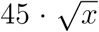.

**Supplementary Figure 3:**
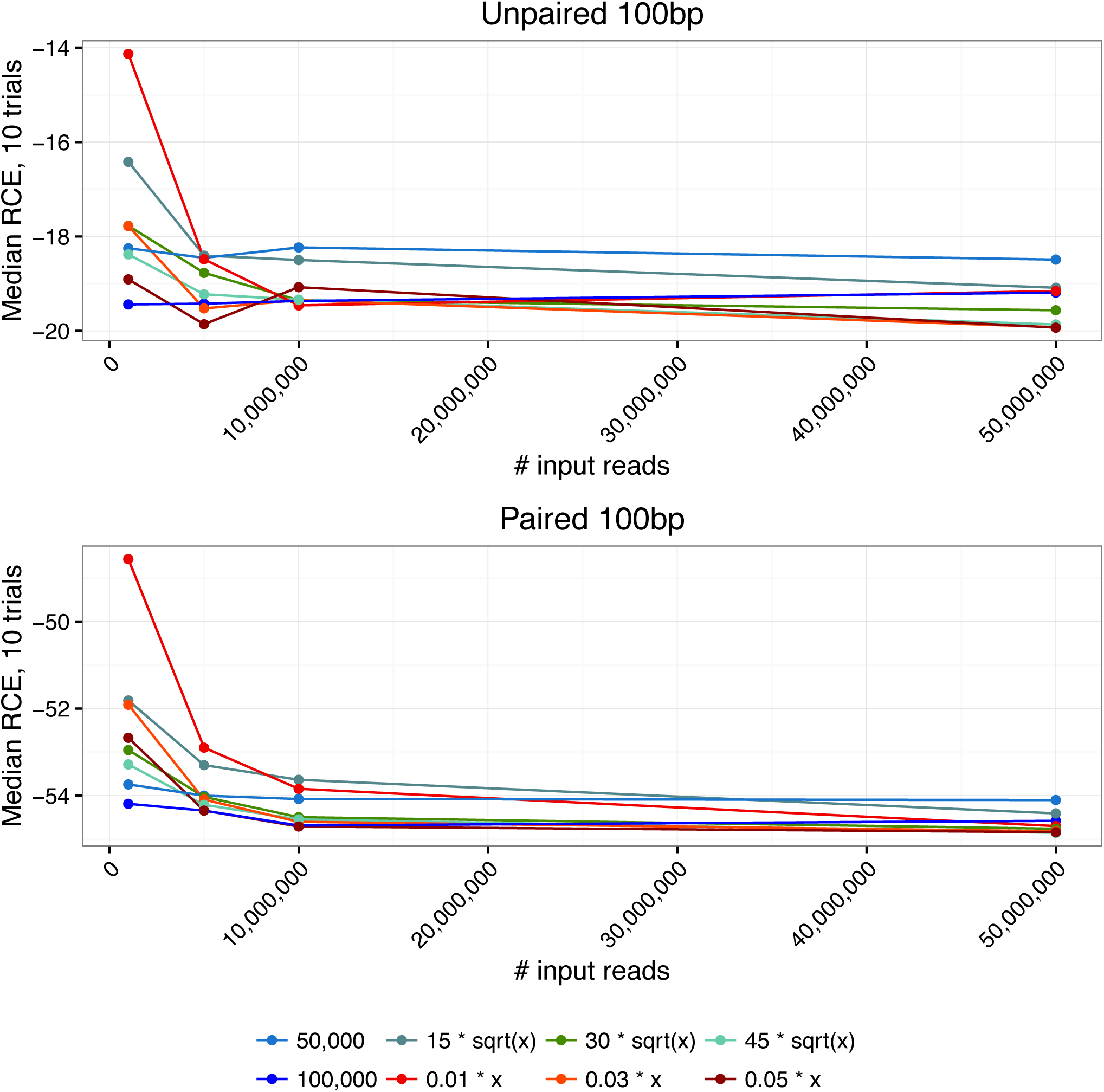
Series of experiments showing how formulas for setting the target number of tandem alignments affect accuracy, measured as the median relative change in squared error across 10 trials.

## Supplementary Note 11 Input modeling and tandem simulation

### Scanning the input

The construct the input model, Qtip using reservoir sampling to obtain an appropriate-sized subset of the input alignments. It does this over the course of a single scan through the input alignments. For paired-end alignments, Qtip’s scan matches up the two ends so that they can be considered together here. Reads that fail to align are ignored, as are non-primary alignments,. During the scan, Qtip distinguishes between the four different categories of alignment: *conc*, *disc*, *unp* and *bad-end*. A separate reservoir sampler is used for each category. The maximum number of alignments sampled in a category can be configured with Qtip’s `--input-model-size` parameter (default: 30,000).

### Templates

Every category of template contains at least the following fields:

- Score
- Read length
- Strand
- Quality string
- Edit transcript

Alignment score is copied from the `AS:i` SAM field. Read length is easily calculated from the SAM/BAM `SEQ` field. Strand is inferred from the SAM `FLAG` field. Quality string is copied from the SAM `QUAL` field.

Edit transcript is inferred from a different combination of SAM fields depending on the aligner used. For Bowtie 2 and BWA-MEM, it is inferred from the `CIGAR` and `MD:Z` fields. SNAP does not report the `MD:Z` field, and the CIGAR field alone is not always sufficient to determine where mismatches are located. However, SNAP can optionally print a CIGAR string that does contain sufficient information to determine where mismatches are located (the `-=` parameter), so when running SNAP from Qtip, that parameter is always specified. When that parameter is specified, the edit transcript can be inferred from the CIGAR field alone.

The 5 fields listed in the previous subsection are sufficient for representing an unpaired read template. A bad-end template includes two additional fields:

- Whether aligned end is end 1 or 2
- Length of the (unaligned) opposite end

These are inferred from the SAM `FLAG` and `SEQ` fields for the opposite end.

A *conc* template contains all of the fields listed above plus these additional fields:

- Score of the opposite end
- Sum of scores of the two ends
- Strand of opposite end
- Quality string of opposite end
- Edit transcript of opposite end
- Whether end 1 aligns to the left of end 2 with respect to the reference
- Fragment length

Most of these additional fields are inferred straightforwardly from the SAM records for the two ends as described previously. Which end aligns upstream is determined by examining the SAM `POS` field for the two records. Fragment length is inferred from the SAM `TLEN` field.

A *disc* template is identical to a *conc* template. Note, though, that the fragment length field may not be interpretable for a discordantly aligned pair, since the two discordantly-aligned ends can align very far apart or even to different chromosomes.

### Handling soft clipping

Input alignments with soft-clipped bases (`S` CIGAR operation) pose a problem for our templates: because the soft-clipped bases failed to align, we do not know how to represent them in the edit transcript. Simply excluding those bases from the edit transcript – effectively trimming them from the tandem read – is not desirable. We would like the tandem reads to mimic the input reads in all important ways, including in the prevalence of soft-clipping.

One solution is to, for each instance of soft clipping, extract the corresponding stretch of reference bases and perform post-facto alignment of the soft-clipped bases. The alignment would likely be poor (hence the soft clipping), but would yield informative values for the edit transcript. This is slow in practice, however, since it requires that we extract a arbitrary stretch of the reference genome – requiring disk head movement in many cases – for each instance of soft clipping.

Instead, Qtip represents soft-clipped bases with special `S` characters in the edit transcript. When simulating the read, Qtip simply inserts a random base into each softclipped position. Advantages of this approach are: (a) it is fast and simple, requiring no additional alignment, and (b) random bases are a reasonable proxy for soft-clipped bases since’ they are unlikely to align well to the reference, and so will likely be clipped. The disadvantage is that there are no guarantees about how successfully the random bases will act as a proxy for the soft-clipped bases. For example, the random bases could, by coincidence, match the reference genome well enough that they align without any soft clipping, or with less soft clipping than was needed for the corresponding input read. This causes the tandem reads to depart somewhat from the character of the input reads, though Qtip’s good performance on local read aligners suggests that the effect is likely minor.

### Simulating from templates

Qtip uses the input models to create the tandem read set in the following way. The user specifies the location of the reference genome in FASTA format to Qtip. Qtip iterates through overlapping windows (*chunks*) of the reference genome. Each chunk consists of *B* + *O* contiguous bases, where the base length *B* is 100,000 by default. We discuss later how the overlap length *O* is determined. The *O* bases at the end of the *B* − *O* sized chunk will appear again at the beginning of the following chunk, except in cases where, for example, we reach the end of a chromosome.

Overlaps between chunks are to ensure all possible read starting positions are equally probable. When a read is simulated, the read’s leftmost base must fall within the first *B* bases of the current chunk. The read’s leftmost base may not fall within the final *O* bases (though those bases will appear again at the beginning of the following chunk).

For each chunk and each alignment category, Qtip calculates the number of reads to simulate according to a binomial draw with *n* = *B*, the number of possible starting positions for the fragment, and 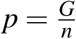 where *G* is the genome length and *n* is the number of reads of that category we aim to simulate in total. *G* is estimated from the FASTA file sizes, and *n* is set according to the `--sim-*` arguments passed to Qtip.

Each unpaired read is then simulated by (1) picking a template of the appropriate type uniformly at random from among those sampled, (2) picking a leftmost read position from among the first *B* positions of the chunk uniformly at random, (3) extracting an appropriate-length substring of the reference genome starting at the chosen position, (4) mutating the extracted reference substring according to the strand and edit transcript information in the template, (5) setting the read name equal to a special string that encodes the read’s true point of origin, (6) setting the read quality string equal to the quality string in the template, and finally (7) printing the read to a FASTQ file in preparation for the tandem alignment step.

A bad-end read is simulated similarly except that a dummy opposite end is synthesized by generating a random nucleotide sequence. The dummy opposite end is virtually guaranteed not to align, mimicking the situation for the input pair.

Paired-end reads are simulated similarly except that when extracting the substring from the reference genome, a substring corresponding to the entire fragment is extracted. In the case of a *conc* pair, fragment length is dictated by the length given in the template. In the case of a *disc* pair, fragment length is set to a pre-determined value larger than the largest concordant fragment length simulated. This is configurable via Qtip’s `--max-allowed-fraglen` parameter. Once a fragment substring is extracted, read sequences are extracted from either end of the fragment, choosing the upstream end and the orientations according to the template.

In order to simulate a long fragment, the entire fragment must be present in some chunk returned by the FASTA parser. To guarantee this will be the case, Qtip sets the overlap length *O* equal to the maximum fragment or read length in any of the input models.

### Aligning tandem reads

After tandem reads are simulated, they are aligned (step 4) using the same read aligner and the same alignment parameters as were used to align the input reads in step 1. However, circumstances can cause the tandem alignments to depart from the template.

First, a tandem read may simply fail to align, decreasing the number of Qtip’s training examples by one. This is relatively rare, since the templates themselves correspond to reads that aligned successfully, and are therefore unlikely to run seriously afoul of the aligner heuristics. However, the tandem read might be extracted from a qualitatively different genomic region than the original read. For example, the tandem read might be drawn from a repeat, whereas the original read was not. This increases the chance that, e.g., the aligner might stop early before finding a valid alignment for the tandem read.

Second, the tandem read might align successfully, but with a different alignment score and/or edit transcript from those in the template. E.g., if the tandem read is drawn from a repetitive portion of the genome, the edits induced by the transcript might cause the read to align with higher alignment score elsewhere. Alternately, the aligner might fail to align the read to its true point of origin and instead align it such that its score is lower than the template score. We so no strong evidence that Qtip’s predictions are systematically biased in either direction, either producing tandem alignments with overall higher or overall lower scores than the input alignments.

### Discussion

We described a method for deriving input models from aligned reads. We also described a method for using the input models, together with the reference genome, to simulate a collection of tandem reads that mimic the input reads in key ways. The method has several advantages: (a) it is simple, not requiring an external read simulator, (b) input models are built during a single pass over the input alignments, (c) reads are simulated during a single pass over the reference genome, (d) since the input alignments are scanned one-or two-at-a-time, and since only a single chunk of the reference genome considered at a time during simulation, the added memory footprint is low.

It should also be noted that, because read sequences are drawn randomly from across the genome and are matched with templates randomly, the tandem reads taken as a whole are not consistent with any particular genome sequence. That is, the reads will not generally agree with each on, e.g., where any SNPs or other variants are located with respect to the reference genome. But since the read aligners themselves consider each read independently and one at a time, this should not affect the performance of the mapping quality model trained in step 6.

## Supplementary Note 12 Feature importances

We studied the feature importances reported by the `RandomForestRegressor` model for the simulation experiments described in the main text in Figure 2 and Table 2. Feature importances are calculated by `scikit-learn` and included in the object representing the fit model. Importances range from 0 to 1, with higher importance indicating the feature tends to appear higher in the decision trees.

Recall that a separate model is trained for each of the four categories of alignment: `conc`, `disc`, `bad-end`, and `unp`. The first three are relevant for the paired-end simulations and the last for unpaired simulations. We plot importances for the 6 features with the highest average importance across all trials and reference genomes. The boxplots show the importances across the 10 trials.

Recall that the random forest model is indifferent to scale. Some of the features described below should intuitively have a positive correlation with mapping quality (e.g. Score diff) while others should have a negative correlation (e.g. % repetitive). The random forest is not predisposed to one or the other.

### Bowtie 2

Supplementary Figure 4 shows results for the Bowtie 2 experiments for 100 nt reads for all reference genomes and models. Supplementary Figure 5 shows the same for 250 nt reads. The features mentioned in those plots are:

- Best score: Alignment score of the reported (best) alignment.
- Score diff: Score of best alignment minus score of second-best alignment, using the configured scoring scheme. If no second-best alignment is found, this is set to a value larger than the largest observed score difference.
- Score diff other: Like Score diff but for the opposite rather than the current end of the paired-end alignment.
- Conc score diff: Score of best concordant paired-end alignment minus score of second-best concordant paired-end alignment, using the configured scoring scheme. The score of a concordant alignment is calculated as the sum of the alignment scores of both ends. If no second-best concordant paired-end alignment is found, this is set to a value larger than the largest observed difference.
- Score diff other: Same as Score diff but for the opposite rather than the current end of the paired-end alignment.
- Edit diff: Edit distance of best alignment minus edit distance of second-best alignment, ignoring the configured scoring scheme. If no second-best alignment is found, this is set to a value larger than the largest observed edit distance difference.
- Edit diff other: Same as Edit diff but for the opposite rather than the current end of the paired-end alignment.
- Conc edit diff: Edit distance of best concordant paired-end alignment minus edit distance of second-best concordant paired-end alignment, ignoring the configured scoring scheme. The edit distance of a concordant alignment is calculated as the sum of the edit distances of both ends. If no second-best concordant paired-end alignment is found, this is set to a value larger than the largest observed difference.
- % unique stranded: Fraction of seeds from the same strand as the reported alignment that matched the reference in exactly one location.
- % unique stranded other: Same as % unique stranded but for the opposite rather than the current end of the paired-end alignment.
- % unique: Fraction of seeds that matched the reference in exactly one location, regardless of seed’s strand.
- % repetitive: Fraction of seeds that matched the reference in more than one location, regardless of seed’s strand.
- Avg seed hits: Average number of times a seed matched the reference.
- Avg seed hits stranded: The average number of times a seed from the same strand as the reported alignment matched the reference.
- Frag length: Inferred fragment length as reported in the SAM `TLEN` field.

### BWA-MEM

Supplementary Figure 6 shows results for the BWA-MEM experiments for 100 nt reads for all reference genomes and models. Supplementary Figure 7 shows the same for 250 nt reads. The features mentioned in those plots are:

- Best score: Alignment score of the reported (best) alignment.
- Score diff: Score of best alignment minus score of second-best alignment, using the configured scoring scheme.
- Score diff o: Like Score diff but for allows the second-best alignment to be redundant with the first.
- Score diff other: Like Score diff but for the opposite rather than the current end of the paired-end alignment.
- Score diff o other: Like Score diff o but for the opposite rather than the current end of the paired-end alignment.
- Conc score diff: Score of best concordant paired-end alignment minus score of second-best alignment, using the configured scoring scheme. The score of a concordant alignment is calculated as the sum of the alignment scores of both ends.
- # tied for 2nd: Number of alignments known to be tied at the second-best alignment score.
- # tied for 2nd o: Like # tied for second but for allows the second-best alignment to be redundant with the first.
- # tied for 2nd o other: Like # tied for second o but for the opposite rather than the current end of the paired-end alignment.
- # conc tied for second: Number of concordant alignments known to be tied at the second-best alignment score. The score of a concordant alignment is calculated as the sum of the alignment scores of both ends.
- Seed coverage: Fraction of read length covered by aligning seeds.
- % repetitive: Fraction of seeds that matched the reference in more than one location, regardless of seed’s strand.
- Frag length: Inferred fragment length as reported in the SAM `TLEN` field.

### SNAP

Supplementary Figure 8 shows results for the SNAP experiments for 100 nt reads for all reference genomes and models. Supplementary Figure 9 shows the same for 250 nt reads. The features mentioned in those plots are:

- Prob best: SNAP’s estimate of the probability the best alignment is correct.
- Prob best other: Like Prob best but for the opposite rather than the current end of the paired-end
- Score diff: Edit distance of best alignment minus edit distance of second-best concordant paired-end alignment, ignoring the configured scoring scheme. (Note: SNAP’s scoring scheme is edit distance, so “edit distance” and “score” are used interchangeably here.) If no second-best alignment was found, this is set to a value just smaller than the most negative observed difference.
- Score diff other: Like Score diff but for the opposite rather than the current end of the paired-end
- Min hits: The minimum number of matches for any seed, regardless of strand.
- Min hits other: Like Min hits other but for the opposite rather than the current end of the paired-end
- Min hits strand: The minimum number of matches for any seed taken from the same strand as the reported alignment.
- Min hits strand other: Like Min hits strand but for the opposite rather than the current end of the paired-end alignment.
- Pop seeds skip: Number of seeds ignored because they matched the reference too many times.
- Hits per lookup: Average number of seed hits per index query.
- Hash misses: Number of index queries that returned 0 seed hits.
- Frag length: Inferred fragment length as reported in the SAM `TLEN` field.

**Supplementary Figure 4:**
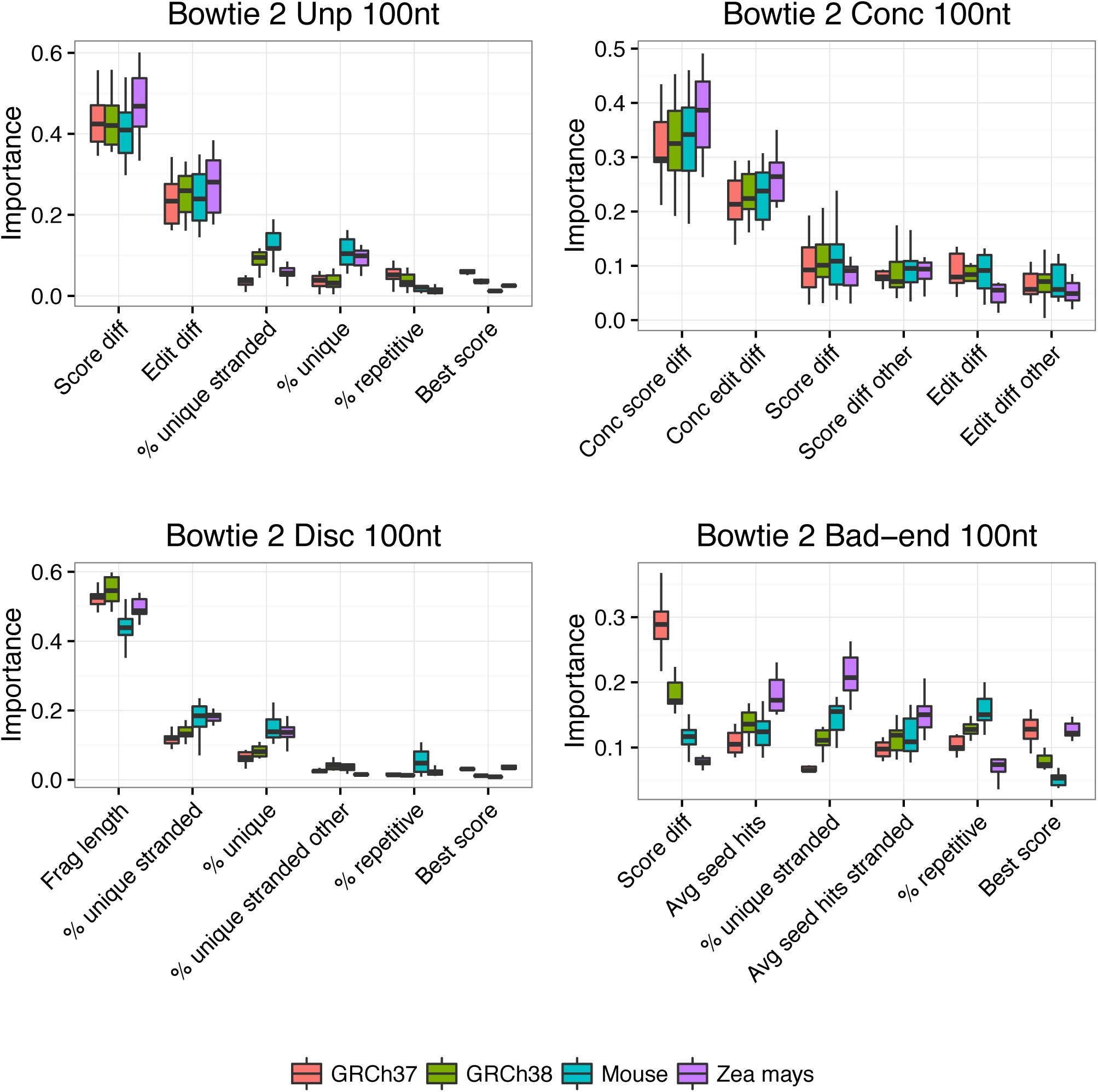
Feature importances for Bowtie 2 experiments on 100 nt reads for all reference genomes and models. Plotted are the 6 features with greatest average importance across all trials and reference genomes. Boxplots show importance measurements across 10 trials. The model for the unpaired simulation is labeled “Unp” and models for the paired-end simulations are labeled “Conc,” “Disc” and “Bad-end.” A more detailed description of the features is given in Supplementary Note 12.

**Supplementary Figure 5:**
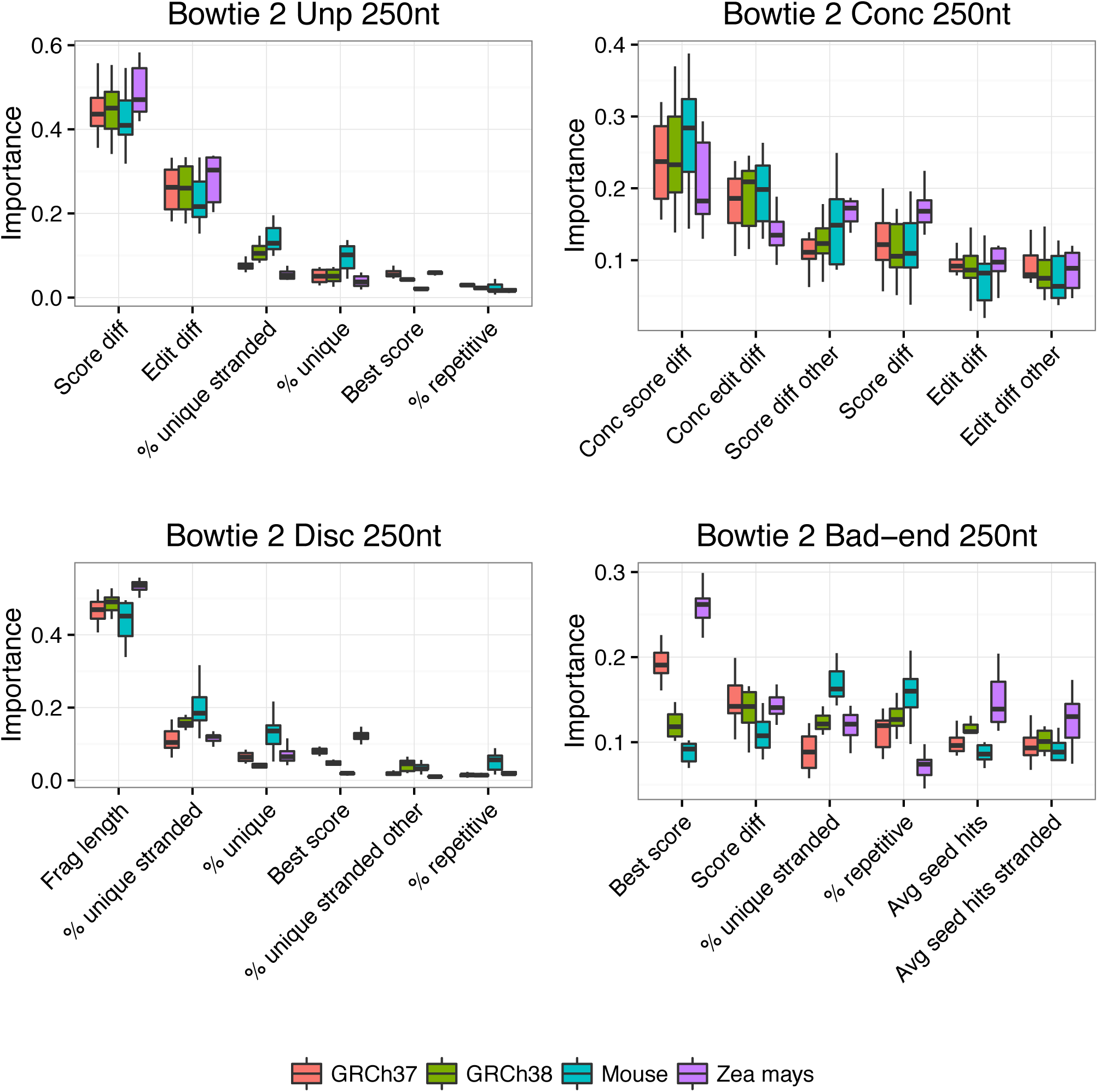
Feature importances for Bowtie 2 experiments on 250 nt reads for all reference genomes and models. Plotted are the 6 features with greatest average importance across all trials and reference genomes. Boxplots show importance measurements across 10 trials. The model for the unpaired simulation is labeled “Unp” and models for the paired-end simulations are labeled “Conc,” “Disc” and “Bad-end.” A more detailed description of the features is given in Supplementary Note 12.

**Supplementary Figure 6:**
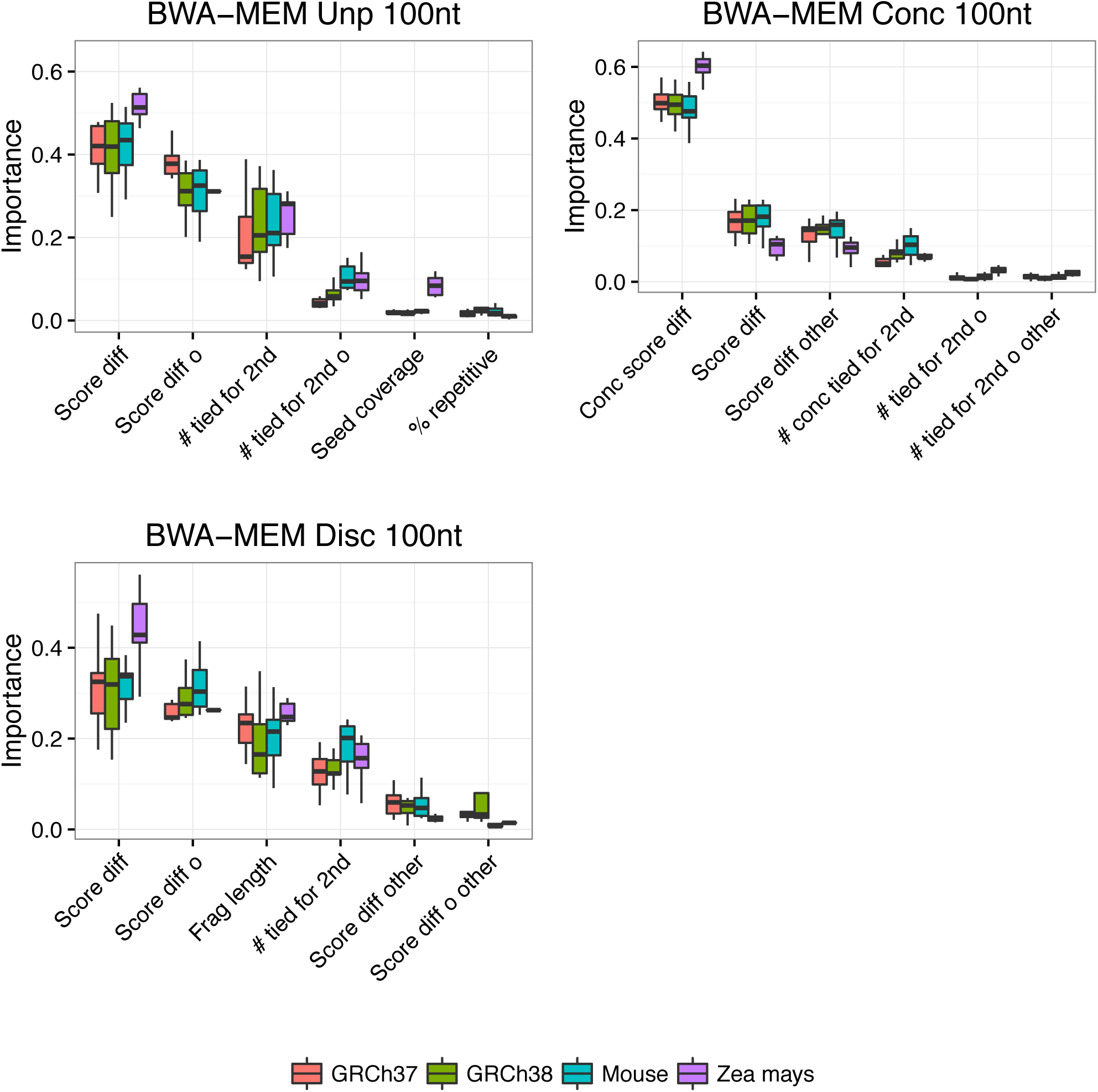
Feature importances for BWA-MEM experiments on 100 nt reads for all reference genomes and models. Plotted are the 6 features with greatest average importance across all trials and reference genomes. Boxplots show importance measurements across 10 trials. The model for the unpaired simulation is labeled “Unp” and models for the paired-end simulations are labeled “Conc” and “Disc.” “Bad-end” is absent because BWA-MEM does not report any instances where only one end of a pair aligns. A more detailed description of the features is given in Supplementary Note 12.

**Supplementary Figure 7:**
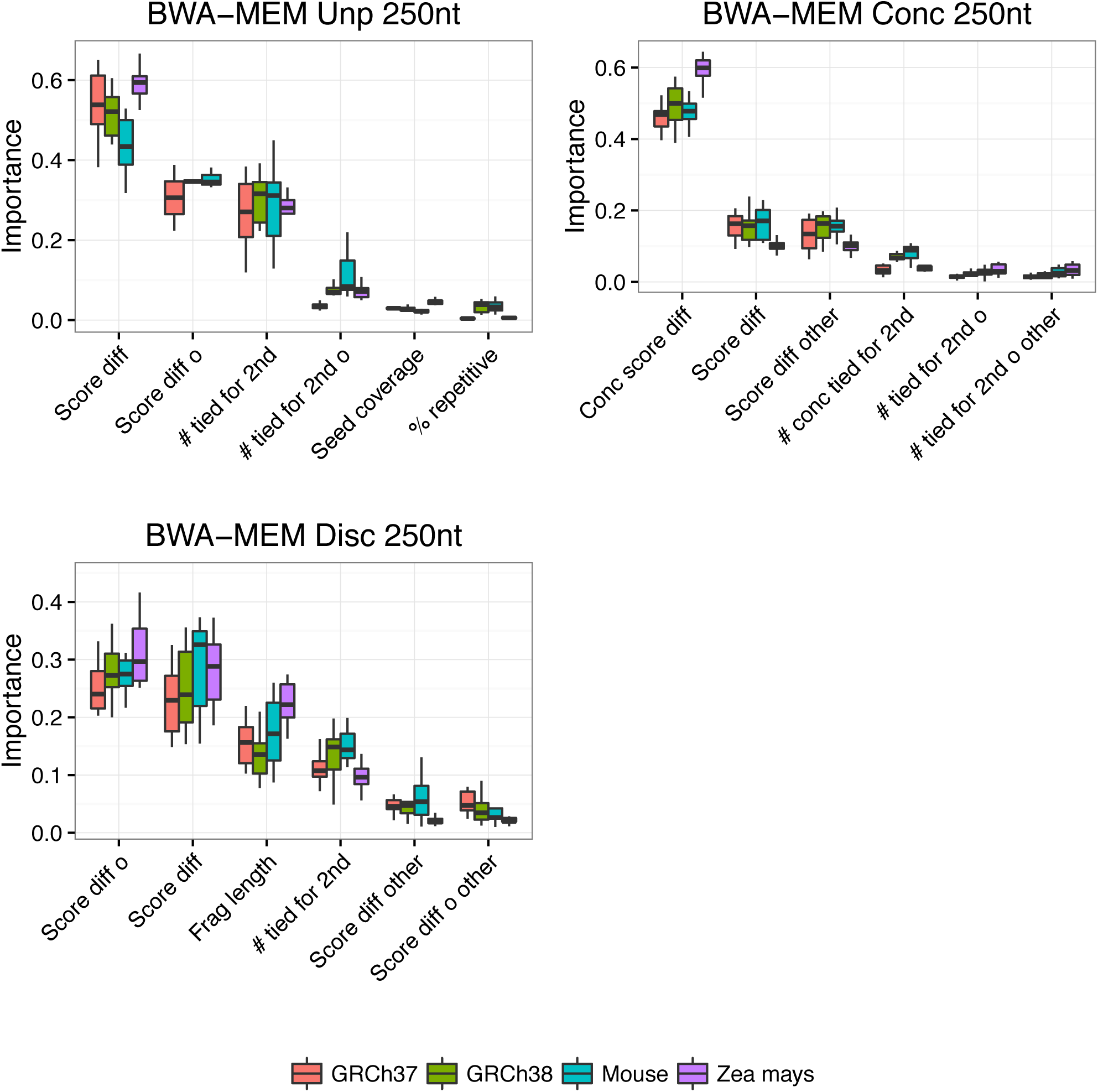
Feature importances for BWA-MEM experiments on 250 nt reads for all reference genomes and models. Plotted are the 6 features with greatest average importance across all trials and reference genomes. Boxplots show importance measurements across 10 trials. The model for the unpaired simulation is labeled “Unp” and models for the paired-end simulations are labeled “Conc” and “Disc.” “Bad-end” is absent because BWA-MEM does not report any instances where only one end of a pair aligns. A more detailed description of the features is given in Supplementary Note 12.

**Supplementary Figure 8:**
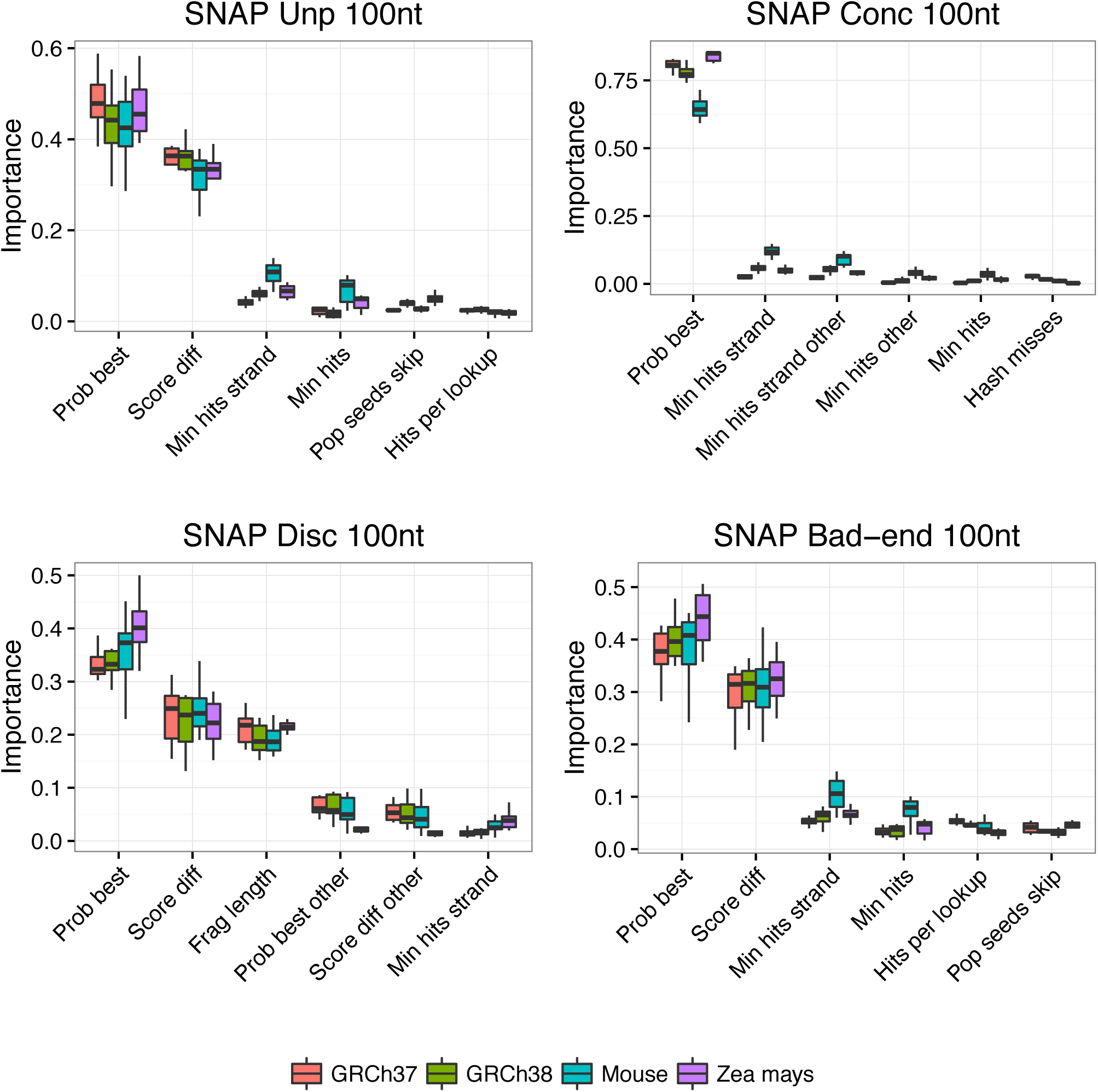
Feature importances for SNAP experiments on 100 nt reads for all reference genomes and models. Plotted are the 6 features with greatest average importance across all trials and reference genomes. Boxplots show importance measurements across 10 trials. The model for the unpaired simulation is labeled “Unp” and models for the paired-end simulations are labeled “Conc,” “Disc” and “Bad-end.” A more detailed description of the features is given in Supplementary Note 12.

**Supplementary Figure 9:**
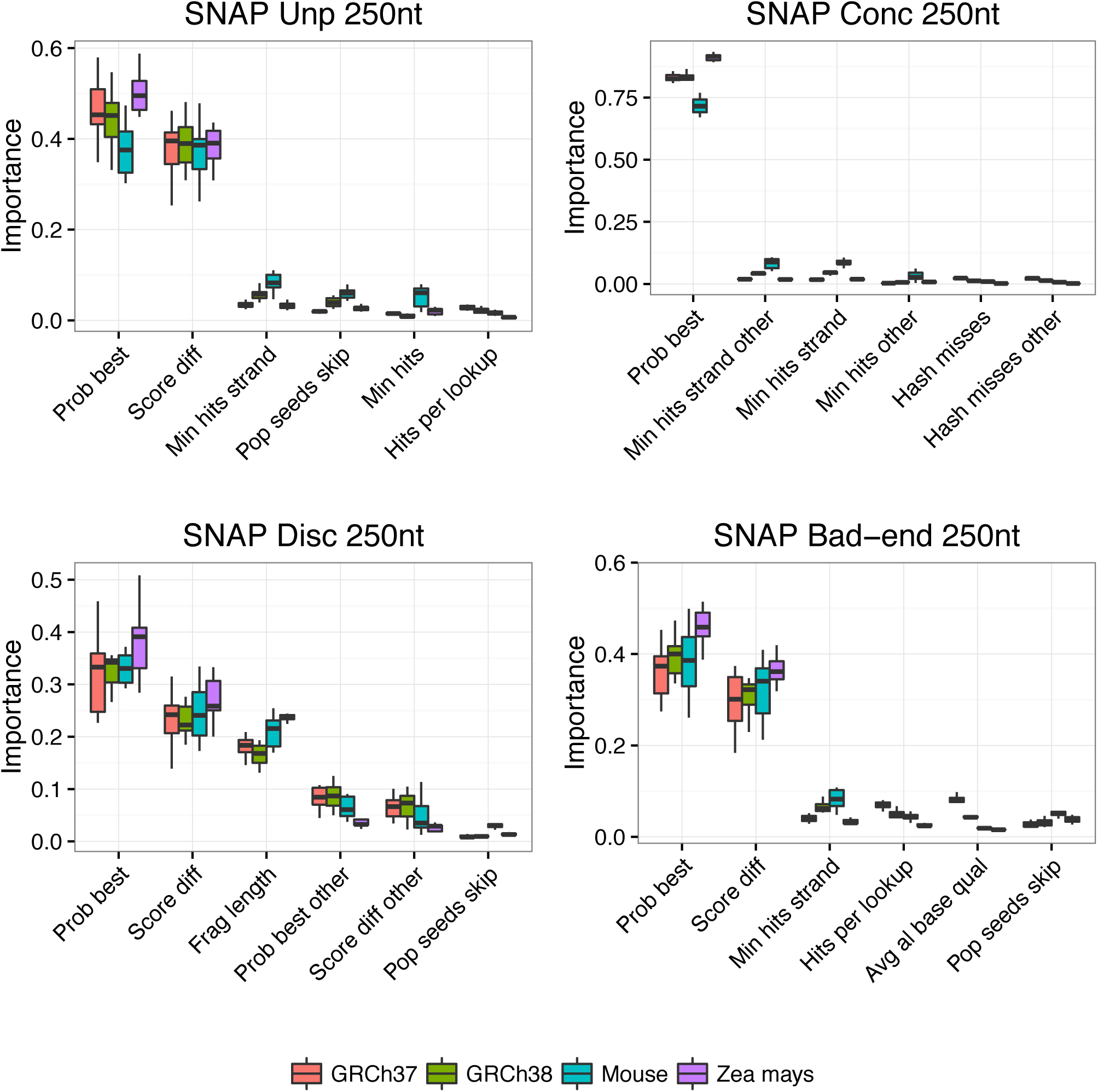
Feature importances for SNAP experiments on 250 nt reads for all reference genomes and models. Plotted are the 6 features with greatest average importance across all trials and reference genomes. Boxplots show importance measurements across 10 trials. The model for the unpaired simulation is labeled “Unp” and models for the paired-end simulations are labeled “Conc,” “Disc” and “Bad-end.” A more detailed description of the features is given in Supplementary Note 12.

## Supplementary Note 13 Mapping-quality prediction model

### Random forest model

Qtip uses the scikit-learn Python module [14] to build its mapping quality prediction model. Specifically Qtip uses scikit-learn’s `RandomForestRegressor`, a model consisting of an ensemble of decision trees. Each tree is trained on a bootstrap sample of the training data, and each tree contributes a “vote” as to the probability the given alignment is correct. The final prediction is an average of all trees’ votes.

Random forests have many advantages in this setting. First, decision tree training is not sensitive to how features are scaled. When adding a new feature, an aligner author can simply report the data as-is, without first having to consider whether log scaling, or any other monotone scaling, is needed to bring the feature into agreement with other features. This is a major advantage over scale-sensitive models like support vector machines or neural networks.

Second, random forests are efficient. They have few hyperparameters, making the training process simpler and faster. E.g., the SNAP aligner takes over 2 hours to align the paired-end ERR050083 dataset, whereas the resulting random forests take about 2 minutes to train. This is a key advantage over, e.g., neural networks. Having said that, the prediction step – step 7 – is the single biggest contributor to Qtip’s computational overhead. We plan to explore whether using a different random forest implementation could increase prediction throughput.

Finally, after a `RandomForestRegressor` model is fit, it optionally reports feature importances. A feature’s importance is essentially the degree to which splitting on the feature helps to separate positive from negative training examples.

### Hyperparameters

Parameters governing the size, shape and splitting behavior of the random forest, often called hyperparameters, have to be chosen at model fitting time. For this model, the relevant parameters are:

- Number of decision trees in the forest. Set to 30 by default in Qtip, but adjustable using the `--num-trees` parameter.
- Maximum number of leaf nodes per decision tree. Set to 35 by default in Qtip, but adjustable using the `--max-leaf-nodes` parameter.
- Maximum number of features to split on at each internal decision tree node. This is expressed as the fraction of the total number of features, and it’s allowed to vary from 0.1 to 0.45 by default. It is adjustable using the `--max-features` parameter.

As mentioned, the `--max-features` hyperparameter is not fixed. In fact, all three hyperparameters can be configured to take values in a range; e.g., specifying the parameter `--num-trees 30,35,40,45,50` will allow `--num-trees` to take on any of the specified values. For each hyperparameter allowed to vary in this way, a particular value is chosen at model-training time via a hyperparameter fitting procedure that performs hill-climbing with a configurable numeric tolerance (`--optimization-tolerance`). The hyperparameters selected are those that yield the best out-of-bag score after training. The out-of-bag score is calculated and by the `RandomForestRegressor` model. As an alternative to the out-of-bag score, cross validation can be used instead (via Qtip’s `--no-oob` parameter).

### Alternate models

Qtip can optionally be configured to use the `ExtraTreesRegressor` class, which implements a variant on random forests called extremely randomized trees [6]. The hyperparameters are defined, configured, and fit in exactly the same manner as for the `RandomForestRegressor`.

### Forming the training and test matrices

The test matrix is a matrix where each row is an input alignment and each column is a feature. Elements contain feature data. In the case of a paired-end alignment, the two ends appear as separate rows in the matrix. The training matrix is similar, except that the rows are tandem alignments rather than input alignments. For each alignment category, Qtip builds a training matrix and uses it to train the model (step 6). Qtip uses the model to predict mapping quality for the input alignments (step 7).

As mentioned, the columns of the matrices are features and the rows are alignments. Qtip might simplify the matrix in certain ways. For example, if all elements in a column are identical in the training matrix, that column is removed from both the training and test matrices. Also, if two columns of the training matrix are identical, the rightmost of the two columns is removed from both the training and test matrices.

To maintain a low memory footprint, matrices are handled such that only a single fixed-sized block of consecutive rows needs to be present in memory at a time. For example, when Qtip parses test records derived from the input SAM file, the records are read in blocks, with only one block residing in memory at a time. By default, a single block contains at most 250K records. This is configurable via Qtip’s `--max-rows` option. Qtip then uses the model to predict mapping qualities for the block and writes the corresponding mapping qualities to disk before deallocating the block and moving on.

### Missing values

As discussed in Supplementary Note 6, some feature data can be “missing.” For example, an important feature used in Qtip is the difference between the score of the best and second-best alignments found by the aligner. But if the aligner fails to find a second-best alignment, this feature will take the value `NA`, meaning missing or “not available.” Qtip deals with `NA` values by automatically setting all of the `NA` values in the column equal to *x* + 1 where *x* is the maximum non-`NA` value in the column. The fact that `NA`s are set to be slightly larger than the largest non-`NA` value often suggests particular way of ordering the data values for a feature. For example, in the case of the difference between the best and second-best score, it makes sense for larger differences to be represented by larger numbers, so that when an `NA` is replaced by *x* + 1, this is interpreted by the model, sensibly, as a large difference.

## Footnotes

1 For a group of alignments sharing the same *Q*, the penalty is averaged across the group’s elements in *C* and *C′*. I.e. if, *â*_*k*_*, â*_*k*+1_*, …, â*_*l*_ is a maximal stretch of alignments sharing the same quality, then 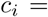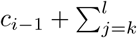 incorrect 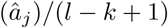 for *k ≤ i ≤ l*.

2 For a group of alignments sharing the same *Q*, the corresponding elements of *E* and *E′* equal the mean squared error of the group. i.e. if, *â*_*k*_*, â*_*k*+1_*, …, â*_*l*_ is a maximal stretch of alignments sharing the same quality, then *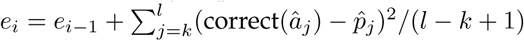* for *k ≤ i ≤ l*.

## References

1. Knut Reinert,Ben Langmead, David Weese, and Dirk J Evers. Alignment of next-generation sequencing reads. Annual review of genomics and human genetics, 16:133–151, 2015.

2. Heng Li, Jue Ruan, and Richard Durbin. Mapping short dna sequencing reads and calling variants using mapping quality scores. Genome research, 18(11):1851–1858, 2008.

3. Heng Li, Bob Handsaker, Alec Wysoker, Tim Fennell, Jue Ruan, Nils Homer, Gabor Marth, Goncalo Abecasis, Richard Durbin, et al. The sequence alignment/map format and samtools. Bioinformatics, 25(16):2078–2079, 2009.

4. Ben Langmead and Steven L Salzberg. Fast gapped-read alignment with bowtie 2.Nature methods, 9(4):357–359, 2012.

5. Heng Li. Aligning sequence reads, clone sequences and assembly contigs with bwa-mem. arXiv preprint arXiv:1303.3997, 2013.

6. Matei Zaharia, William J Bolosky, Kristal Curtis, Armando Fox, David Patterson, Scott Shenker, Ion Stoica, Richard M Karp, and Taylor Sittler. Faster and more accurate sequence alignment with snap. arXiv preprint arXiv:1111.5572, 2011.

7. Joseph K. Pickrell, Yoav Gilad, and Jonathan K. Pritchard. Comment on widespread rna and dna sequence differences in the human transcriptome. Science, 335(6074):1302, 2012.

8. Todd J Treangen and Steven L Salzberg. Repetitive dna and next-generation sequenc-ing: computational challenges and solutions. Nature Reviews Genetics, 13(1):36–46, 2011.

9. Margaret Taub, Doron Lipson, Terence P Speed, et al. Methods for allocating ambiguous short-reads. Communications in Information & Systems, 10(2):69–82, 2010.

10. Todd J Treangen and Steven L Salzberg. Repetitive dna and next-generation sequencing: computational challenges and solutions. Nature Reviews Genetics, 13(1):36–46, 2012.

11. Samuel Karlin and Stephen F Altschul. Methods for assessing the statistical significance of molecular sequence features by using general scoring schemes. Proceedings of the National Academy of Sciences, 87(6):2264–2268, 1990.

12. Heng Li and Richard Durbin. Fast and accurate short read alignment with burrows– wheeler transform. Bioinformatics, 25(14):1754–1760, 2009.

13. Heng Li and Richard Durbin. Fast and accurate long-read alignment with burrows– wheeler transform. Bioinformatics, 26(5):589–595, 2010.

14. Sven H Giese, Franziska Zickmann, and Bernhard Y Renard. Specificity control for read alignments using an artificial reference genome-guided false discovery rate. Bioinformatics, 30(1):9–16, 2014.

15. Matthew Ruffalo, Mehmet Koyutϋrk, Soumya Ray, and Thomas LaFramboise. Accurate estimation of short read mapping quality for next-generation genome sequencing. Bioinformatics, 28(18):i349–i355, 2012.

16. Wan-Ping Lee, Michael P Stromberg, Alistair Ward, Chip Stewart, Erik P Garrison, and Gabor T Marth. Mosaik: A hash-based algorithm for accurate next-generation sequencing short-read mapping. PloS one, 9(3):e90581, 2014.

17. Alan Hodgkinson, Jean-Christophe Grenier, Elias Gbeha, and Philip Awadalla. A haplotype-based normalization technique for the analysis and detection of allele specific expression. BMC bioinformatics, 17(1):364, 2016.

18. Manuel Holtgrewe. Mason–a read simulator for second generation sequencing data. Technical Report FU Berlin, 2010.

19. Erik Garrison and Gabor Marth. Haplotype-based variant detection from short-read sequencing. arXiv preprint arXiv:1207.3907, 2012.

20. Aaron McKenna, Matthew Hanna, Eric Banks, Andrey Sivachenko, Kristian Cibulskis, Andrew Kernytsky, Kiran Garimella, David Altshuler, Stacey Gabriel, Mark Daly, et al. The genome analysis toolkit: a mapreduce framework for analyzing next-generation dna sequencing data. Genome research, 20(9):1297–1303, 2010.

21. Shin Lin, Benilton Carvalho, David J Cutler, Dan E Arking, Aravinda Chakravarti, and Rafael A Irizarry. Validation and extension of an empirical bayes method for snp calling on affymetrix microarrays. Genome biology, 9(4):R63, 2008.

22. P Green AFA Smit, R Hubley. Repeatmasker open-4.0, accessed feb 4, 2017.

23. P. S. Schnable, D. Ware, R. S. Fulton, J. C. Stein, F. Wei, S. Pasternak, C. Liang, Zhang, L. Fulton, T. A. Graves, P. Minx, A. D. Reily, L. Courtney, S. S. Kruchowski, C. Tomlinson, C. Strong, K. Delehaunty, C. Fronick, B. Courtney, S. M. Rock, E. Belter, F. Du, K. Kim, R. M. Abbott, M. Cotton, A. Levy, P. Marchetto, Ochoa, S. M. Jackson, B. Gillam, W. Chen, L. Yan, J. Higginbotham, M. Cardenas, J. Waligorski, E. Applebaum, L. Phelps, J. Falcone, K. Kanchi, T. Thane, A. Scimone, N. Thane, J. Henke, T. Wang, J. Ruppert, N. Shah, K. Rotter, J. Hodges, E. Ingenthron, M. Cordes, S. Kohlberg, J. Sgro, B. Delgado, K. Mead, A. Chinwalla, S. Leonard, K. Crouse, K. Collura, D. Kudrna, J. Currie, R. He, A. Angelova, S. Ra-jasekar, T. Mueller, R. Lomeli, G. Scara, A. Ko, K. Delaney, M. Wissotski, G. Lopez, D. Campos, M. Braidotti, E. Ashley, W. Golser, H. Kim, S. Lee, J. Lin, Z. Dujmic, W. Kim, J. Talag, A. Zuccolo, C. Fan, A. Sebastian, M. Kramer, L. Spiegel, L. Nascimento, T. Zutavern, B. Miller, C. Ambroise, S. Muller, W. Spooner, A. Narechania, L. Ren, S. Wei, S. Kumari, B. Faga, M. J. Levy, L. McMahan, P. Van Buren, M. W. Vaughn, K. Ying, C. T. Yeh, S. J. Emrich, Y. Jia, A. Kalyanaraman, A. P. Hsia, W. B. Barbazuk, R. S. Baucom, T. P. Brutnell, N. C. Carpita, C. Chaparro, J. M. Chia, J. M. Deragon, J. C. Estill, Y. Fu, J. A. Jeddeloh, Y. Han, H. Lee, P. Li, D. R. Lisch, S. Liu, Z. Liu, D. H. Nagel, M. C. McCann, P. SanMiguel, A. M. Myers, D. Nettleton, J. Nguyen, B. W. Penning, L. Ponnala, K. L. Schneider, D. C. Schwartz, A. Sharma, C. Soderlund, N. M. Springer, Q. Sun, H. Wang, M. Waterman, R. Westerman, T. K. Wolfgruber, L. Yang, Y. Yu, L. Zhang, S. Zhou, Q. Zhu, J. L. Bennetzen, R. K. Dawe, J. Jiang, N. Jiang, G. G. Presting, S. R. Wessler, S. Aluru, R. A. Martienssen, S. W. Clifton, W. R. McCombie, R. A. Wing, and R. K. Wilson. The B73 maize genome: complexity, diversity, and dynamics. Science, 326(5956):1112–1115, Nov 2009.

24. M. Nattestad and M. C. Schatz. Assemblytics: a web analytics tool for the detection of variants from an assembly. Bioinformatics, 32(19):3021–3023, Oct 2016.

25. Michael A Eberle, Epameinondas Fritzilas, Peter Krusche, Morten Källberg, Benjamin L Moore, Mitchell A Bekritsky, Zamin Iqbal, Han-Yu Chuang, Sean J Humphray, Aaron L Halpern, et al. A reference data set of 5.4 million phased human variants validated by genetic inheritance from sequencing a three-generation 17-member pedigree. Genome Research, 27(1):157–164, 2017.

26. Heng Li. Toward better understanding of artifacts in variant calling from highcoverage samples. Bioinformatics, 30(20):2843–2851, Oct 2014.

27. Leo Breiman. Random forests. Machine learning, 45(1):5–32, 2001.

28. F. Pedregosa, G. Varoquaux, A. Gramfort, V. Michel, B. Thirion, O. Grisel, M. Blondel, P. Prettenhofer, R. Weiss, V. Dubourg, J. Vanderplas, A. Passos, D. Cournapeau, M. Brucher, M. Perrot, and E. Duchesnay. Scikit-learn: Machine learning in Python. Journal of Machine Learning Research, 12:2825–2830, 2011.

29. Cole Trapnell, Lior Pachter, and Steven L Salzberg. Tophat: discovering splice junctions with rna-seq. Bioinformatics, 25(9):1105–1111, 2009.

30. Alexander Dobin, Carrie A Davis, Felix Schlesinger, Jorg Drenkow, Chris Zaleski, Sonali Jha, Philippe Batut, Mark Chaisson, and Thomas R Gingeras. Star: ultrafast universal rna-seq aligner. Bioinformatics, 29(1):15–21, 2013.

## References

1. D. A. Benson, M. Cavanaugh, K. Clark, I. Karsch-Mizrachi, D. J. Lipman, J. Ostell, and E. W. Sayers. GenBank. Nucleic Acids Res., 45(D1):D37–D42, Jan 2017.

2. K. Berlin, S. Koren, C. S. Chin, J. P. Drake, J. M. Landolin, and A. M. Phillippy. Assembling large genomes with single-molecule sequencing and locality-sensitive hashing. Nat. Biotechnol., 33(6):623–630, Jun 2015.

3. D. M. Church, V. A. Schneider, K. M. Steinberg, M. C. Schatz, A. R. Quinlan, C. S. Chin, P. A. Kitts, B. Aken, G. T. Marth, M. M. Hoffman, J. Herrero, M. L. Mendoza, R. Durbin, and P. Flicek. Extending reference assembly models. Genome Biol., 16:13, Jan 2015.

4. Michael A Eberle, Epameinondas Fritzilas, Peter Krusche, Morten Källberg, Benjamin L Moore, Mitchell A Bekritsky, Zamin Iqbal, Han-Yu Chuang, Sean J Humphray, Aaron L Halpern, et al. A reference data set of 5.4 million phased human variants validated by genetic inheritance from sequencing a three-generation 17-member pedigree. Genome Research, 27(1):157–164, 2017.

5. Otto Erlwein, Mark J Robinson, Simon Dustan, Jonathan Weber, Steve Kaye, and Myra O McClure. Dna extraction columns contaminated with murine sequences. PLoS One, 6(8):e23484, 2011.

6. Pierre Geurts, Damien Ernst, and Louis Wehenkel. Extremely randomized trees. Machine learning, 63(1):3–42, 2006.

7. Manuel Holtgrewe. Mason–a read simulator for second generation sequencing data. Technical Report FU Berlin, 2010.

8. Weichun Huang, Leping Li, Jason R Myers, and Gabor T Marth. Art: a nextgeneration sequencing read simulator. Bioinformatics, 28(4):593–594, 2012.

9. S. Kurtz, A. Phillippy, A. L. Delcher, M. Smoot, M. Shumway, C. Antonescu, and S. L. Salzberg. Versatile and open software for comparing large genomes. Genome Biol., 5(2):R12, 2004.

10. Heng Li. Toward better understanding of artifacts in variant calling from high-coverage samples. Bioinformatics, 30(20):2843–2851, Oct 2014.

11. Richard W Lusk. Diverse and widespread contamination evident in the unmapped depths of high throughput sequencing data. PloS one, 9(10):e110808, 2014.

12. M. Nattestad and M. C. Schatz. Assemblytics: a web analytics tool for the detection of variants from an assembly. Bioinformatics, 32(19):3021–3023, Oct 2016.

13. Anthony O Olarerin-George and John B Hogenesch. Assessing the prevalence of mycoplasma contamination in cell culture via a survey of ncbi’s rna-seq archive. Nucleic acids research, 43(5):2535–2542, 2015.

14. F. Pedregosa, G. Varoquaux, A. Gramfort, V. Michel, B. Thirion, O. Grisel, M. Blondel, P. Prettenhofer, R. Weiss, V. Dubourg, J. Vanderplas, A. Passos, D. Cournapeau, M. Brucher, M. Perrot, and E. Duchesnay. Scikit-learn: Machine learning in Python. Journal of Machine Learning Research, 12:2825–2830, 2011.

15. Artem Tarasov, Albert J Vilella, Edwin Cuppen, Isaac J Nijman, and Pjotr Prins. Sambamba: fast processing of ngs alignment formats. Bioinformatics, 31(12):2032–2034, 2015.

